# DIST: Distance-based Inference of Species Trees

**DOI:** 10.1101/2025.05.02.651899

**Authors:** Menno J. de Jong, Axel Janke

**Affiliations:** Senckenberg Biodiversity and Climate Research Institute (SBiK-F), Georg-Voigt-Strasse 14-16, Frankfurt am Main, 60325, Germany; Institute for Ecology, Evolution and Diversity, Goethe University, Max-von-Laue-Strasse. 9, Frankfurt am Main, Germany; LOEWE-Centre for Translational Biodiversity Genomics (TBG), Senckenberg Nature Research Society, Georg-Voigt-Strasse 14-16, Frankfurt am Main, Germany

## Abstract

Inferring species trees from concatenated loci is often criticised for failing to account for gene tree discordance – particularly when using character-based methods. However, this criticism does not apply to distance-based concatenation trees, which can be shown to be statistically consistent even in anomaly zones. Building on this insight, we introduce DIST (Distance-based Inference of Species Trees), an intuitive and scalable method that infers species trees from population-level distance matrices containing multi-locus estimates of *D_xy_, F_ST_* or coalescence units (*τ*). DIST derives these values from between-individual sequence dissimilarity estimates, *E(p)*, using basic equations from coalescence theory. Under certain conditions, DIST can also quantify gene tree discordance and distinguish whether it arises from gene flow or incomplete lineage sorting alone. While conceptually related to more sophisticated summary methods, DIST differs in that it does not seek the species tree which best explains a set of gene trees. Instead, it searches for the species tree which best explains an *average* gene tree, of which all branch lengths reflect mean coalescence time, *E(t)*. Although this average gene tree is rarely observed empirically, it is approximated by an individual-level distance-based tree, traditionally referred to as a ‘tree of individuals’. The DIST algorithm is implemented in the R package *SambaR*, which now accepts input in the form of pairwise *E*(*p)* estimates.

## INTRODUCTION

Species trees inferred from multi-locus datasets are state-of-the-art reproductions of Darwin’s Tree of Life. Unlike their single-locus precursors, these genetic trees can account for incomplete lineage sorting and even, by incorporating gene flow edges, for reticulate evolution. These innovations epitomise the remarkable reinvention of phylogenetics in the postgenomic era. With multi-locus data replacing single-locus alignments, the field eagerly transitioned from reconstructing the history of molecules to its foundational goal of reconstructing the history of populations. In less than two decades, this goal has been realised through a wide array of ingenious approaches.

As an alternative to these sophisticated but often complex and computationally expensive approaches, we present a more intuitive and scalable method, called DIST, an acronym for Distance-based Inference of Species Trees. This method simply infers the species tree from population-level distance matrices containing multi-locus *D_xy_, F_ST_*- or *τ*-estimates, calculated from datasets of concatenated loci (Fig. 1). The branch lengths of the resulting species trees either denote mean coalescence time (*D_xy_*), the relative difference in coalescence time within versus between populations (*F_ST_*), or coalescent units (*τ*, i.e., population split time in multiples of population size). Apart from the species tree, DIST also infers individual-level trees, traditionally known as a ‘tree of individuals’ (Mountain and Cavalli-Sforza 1997).

**Figure 1.**
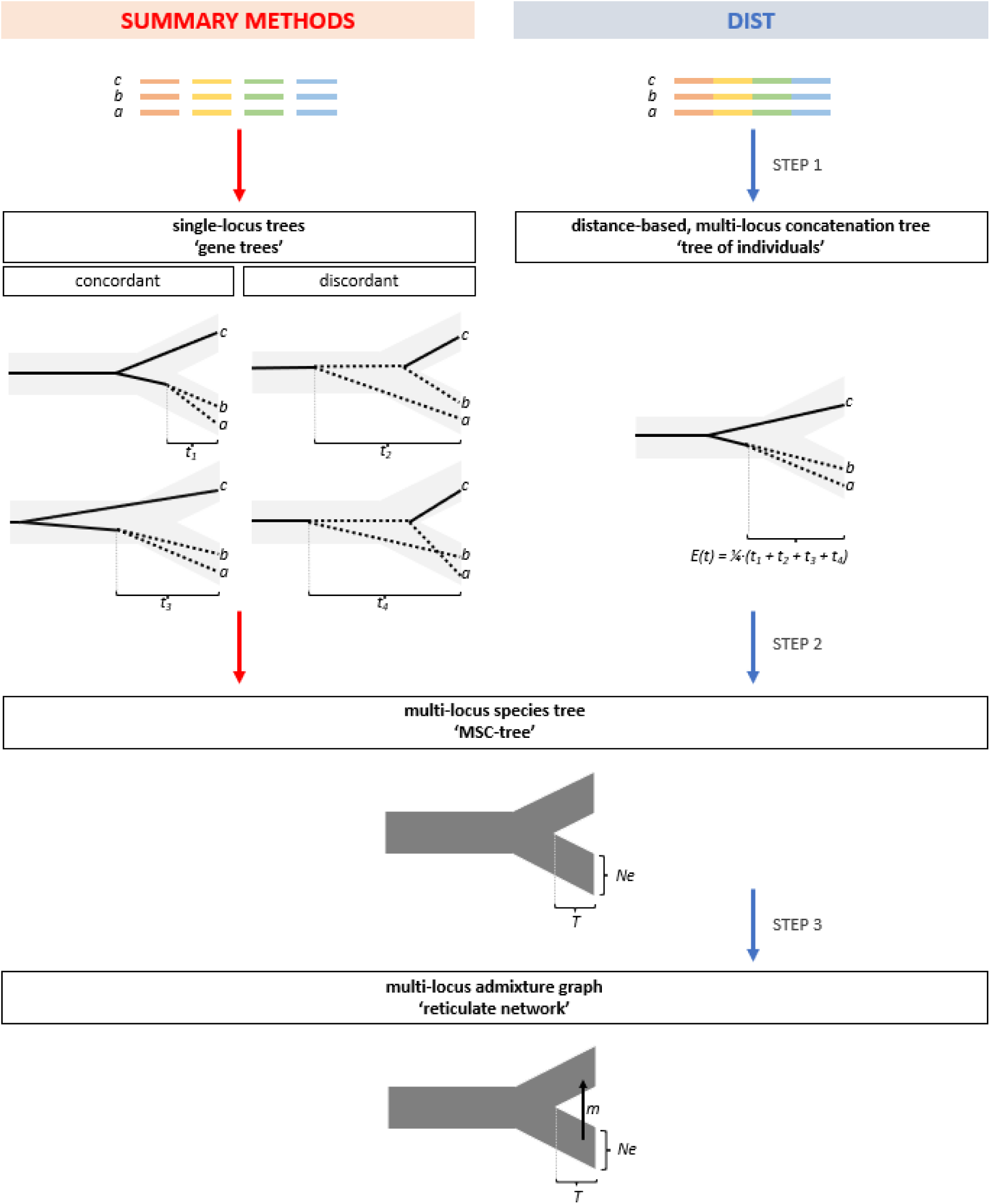
DIST workflow. Coalescent-based summary methods (left) infer the species tree from a set of gene trees. In contrast, DIST (right) infers the species tree from an individual-level concatenation tree, known as a ‘tree of individuals’. The underlying assumption is that this tree of individuals approximates an imaginary average gene tree, of which all branch lengths reflect mean coalescence time. In the optional third step, gene flow edges are added to account for reticulate evolution, thereby transforming the species tree into an admixture graph, also known are reticulate network.

We recognise that many phylogeneticists consider analysing datasets of concatenated loci to be poor practice. The primary concern is the existence of anomaly zones, in which the topology of the most common gene tree differs from the topology of the species tree (Pamilo and Nei 1988; Degnan and Rosenberg 2006; Kubatko and Degnan 2007; Degnan and Rosenberg 2009; Huang and Knowles 2009; Roch and Steel 2015). This does indeed lead to inconsistency if a concatenation tree is equated with the most common gene tree, a procedure known as the ‘democratic vote’ principle (Pamilo and Nei 1988) (Fig. 2).

**Figure 2.**
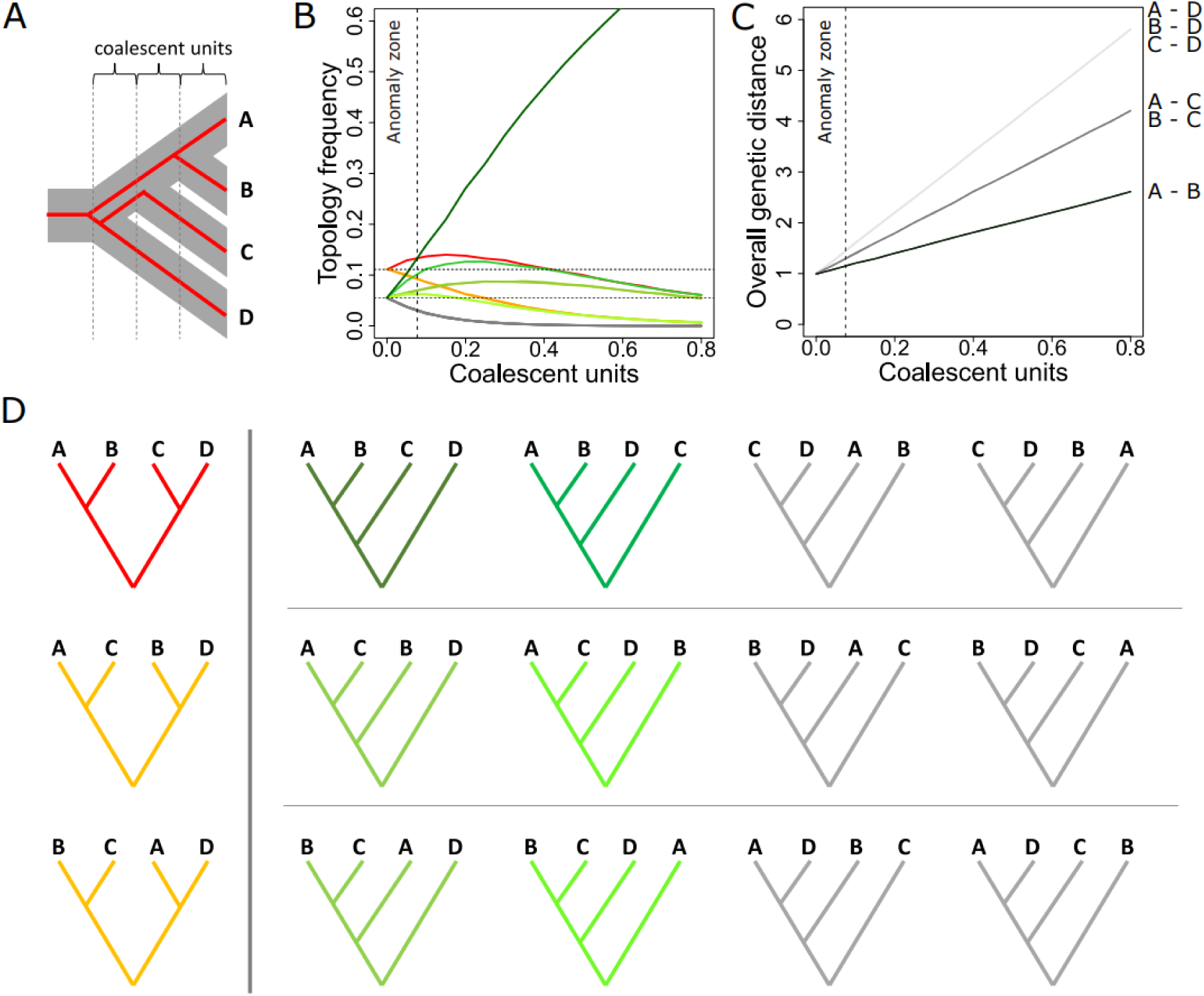
Concatenation also works in the anomaly zone. **A**. Species tree (grey) used for simulating gene trees, together with one particular gene tree (red). **B**. Frequency of topologies of 100.000 simulated gene tree, simulated using the function ‘sim.coaltree.sp’ of the R package ‘phybase’, given the species tree depicted in A, with increasing branch lengths in coalescent units (*τ*, multiples of 2*Ne*), and assuming population size constancy. Colours refer to D. Note that in a panmictic population (*τ* = 0), the three balanced gene trees and the twelve unbalanced gene tree initially occur with a frequency of 1/9 and 1/18, respectively. As a result, when the species tree has short branch lengths (*τ* < 0.08), the most common gene tree is discordant. This is known as the anomaly zone. **C**. A concatenation tree is constructed by calculating a weighted mean of coalescence time, here shown in coalescent units, averaged over all gene trees. Note that even in the anomaly zone the order of genetic distances corresponds with the species tree. **D**. All possible gene trees, coloured according to B, and divided in four subsets for which it is obvious that the four lineages are equidistant if occurring with equal frequencies.

In practice, however, the topology of concatenation trees does not necessarily mirror that of the most common gene tree, particularly not when constructed using a distance-based method. As we confirmed through gene tree simulations with the R package ‘phybase’ (Liu and Yu 2010), multi-locus distances between individuals are weighted averages across all observed gene trees, rather than representing only the most common gene tree (Fig. 2).

When measured in terms of sequence dissimilarity, *E(p)*, genome-wide distances between individuals, and hence mean values between populations (*D_xy_*), are proportional to mean coalescence time, *E(t)*, averaged over all loci. Whether within or outside anomaly zones, the order of *E(t)* among population pairs is expected to mirror the order of population splits, *T* (Criscuolo et al. 2006; A.W.F. Edwards 2009; Liu and Edwards 2009; Dasarathy et al. 2015; ALLMAN et al. 2019). It can also been shown that clustering algorithms based on least squares or minimum evolution principles produce bifurcating trees that accurately represent distance matrices (Pardi and Gascuel 2012). These findings imply that distance-based concatenation trees, whether individual-level or population-level, are statistically consistent.

A common misconception about distance-based analyses is that compressing genomic datasets into a distance matrix with point estimates necessarily results in losing information about incomplete lineage sorting and introgression. In reality, a distance matrix with *E(p)*-estimates contains hidden information about the extent of gene tree discordance within genomes, and even whether this discordance is caused by deep coalescence (i.e., incomplete lineage sorting) or by gene flow instead. DIST extracts this information to construct a distance matrix containing *τ*-estimates and to detect gene flow events, which then serves to infer the species tree and optionally a reticulate network. The underlying algorithms are based on the multispecies coalescent model.

Combining concatenation and coalescence-based approaches into a single method, as implemented in DIST, may appear to some like combining water and fire. The dispute between advocates of the two approaches is long-standing, especially when seen as a continuation of the ‘supermatrix versus supertree’ debate, also known as ‘total evidence versus consensus’ debate (Gadagkar et al. 2005). However, recent advances in the field of phylogenetics have overturned this debate. Concatenation and coalescence-based tree reconstruction, once rivalling approaches, now complement each other.

A modern multispecies coalescent-based supertree represent the species tree. The term ‘summary method’ has effectively become a misnomer, because a species tree does not only summarise gene trees but actually explains them. To this end, supertrees have adopted a novel type of branch lengths, known as drift or coalescence units, *τ*, which denote the product of effective population size (*Ne*) and population split time (*T*). Meanwhile, concatenation trees have remained unchanged. As before, they summarise gene trees by depicting an imaginary average gene tree, with branch lengths representing mean coalescence time, *E(t)*. DIST makes use of the distinction between the two tree types by inferring the population-level distance matrix underlying the species tree from the individual-level distance matrix underlying the tree of individuals.

Despite its simplicity, the rationale behind DIST is related to that of the more sophisticated coalescent-based summary methods. Namely, it assumes that the species tree predicts a probability distribution of gene trees, and that the inverse problem – inferring the species from the gene trees – can be solved using a closed form expression (Liu, Wu, et al. 2015). The key difference lies in how the inverse problem is solved. Whereas summary methods draw inferences from the entire distribution of gene trees (for instance by searching for the species tree that maximises the number of induced quartet trees shared with all gene trees), DIST considers only the imaginary *average* gene tree (by searching for the species tree that best explains the average gene tree).

Given the infinite number of potential combinations of branch lengths for any given set of individuals, this average gene tree will rarely, if ever, be observed in reality. Even when ignoring branch lengths and focusing on topologies alone, the probability that a gene tree exists which is fully concordant with the species tree remains minuscule when sampling a large number of individuals from recently diverged populations. However, while rarely observed empirically, the average gene tree can be approximated by reconstructing the tree of individuals because, by the law of large numbers, concatenation of unlinked loci cancels out single-locus stochasticity.

An advantage of DIST compared to summary methods is that it removes the need to detect unlinked, recombination-free loci. In practice, such elusive ‘c-loci’ are difficult to delineate, and often so small that they contain insufficient informative sites for reliable phylogenetic inference. Most researchers therefore opt to infer gene trees from fixed-sized, arbitrarily delineated windows, which are composites of true c-loci. This violation introduces gene tree estimation errors and, consequently, the risk of flawed species tree inference (Degnan and Rosenberg 2009; Gatesy and Springer 2014; Mailund et al. 2014; Chou et al. 2015; Edwards et al. 2016; Springer and Gatesy 2016; Springer and Gatesy 2018; Doyle 2022). Whereas site-based coalescent approaches bypass c-loci detection by operating on single nucleotides (Chifman and Kubatko 2014; Zhang et al. 2025), DIST takes the exact opposite approach by operating on entire genomes.

Several practical considerations favour DIST over existing species tree inference methods. Its intuitive design and straightforward application, along with built-in validation check points, leave little room for user error. Being distance-based, DIST is computationally inexpensive, so much so that it can process full genome-scale datasets of hundreds of individuals within hours (Subramanian et al. 2019; Xu et al. 2025). This allows for sensitivity analyses and green computing (Kumar 2022). Furthermore, because distances are calculated in a pairwise manner, DIST is robust to the magnitude and variation in levels of missing data across samples. Lastly, unlike species tree inference methods which analyse allele frequency data and therefore require large sample sizes, reliable inferences can, in theory, be made from a single individual per population.

DIST is implemented in the R package SambaR (de Jong et al. 2021), which now accepts, apart from SNP datasets, also input data frames containing pairwise *E(p)*-estimates. A Unix script is provided to generate this data frame from an input gVCF-file, although other tools may be used as well (Subramanian et al. 2019; Xu et al. 2025). Here, we present the DIST algorithm in full detail, starting with providing a historical context.

## 1. THEORETICAL UNDERPINNINGS OF DIST

### A short history of species tree inference

Phylogenetics and population-genetics are intertwined research fields with many shared interests. Preventing their unification is a fundamental, hard-coded incompatibility. A population-geneticist thinks in terms of allele frequencies per population, while a phylogeneticist thinks in terms of population frequencies per allele. Since these two types of conditional probabilities can only be inverted using Bayes rule, the fields have diverged into two parallel yet complimentary paradigms, each with its own specialised language – essentially represent two sides of the same coin (Fig. 3, S1).

**Figure 3.**
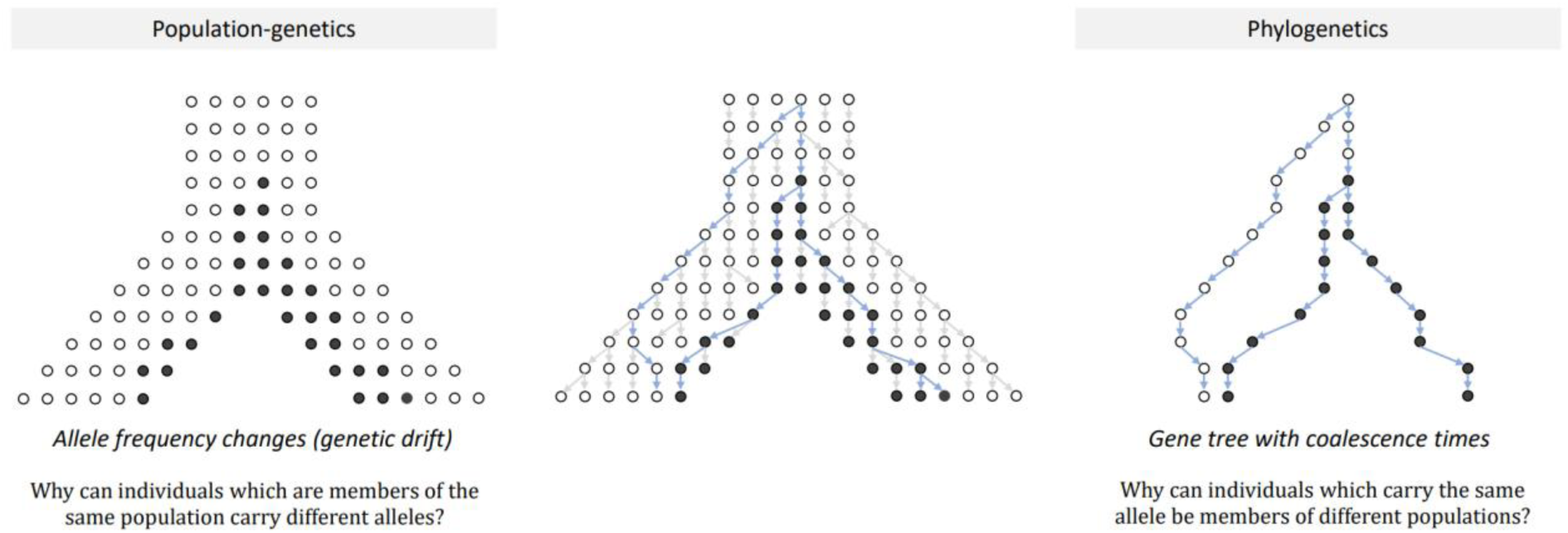
Two sides of the same coin: population-genetics versus phylogenetics. Whereas population-genetics categorise for each locus the observed genetic variation among individuals by population, phylogenetics do so by allele. This has resulted in two major types of species tree inference methods, with species trees explaining either observed allele frequencies or alternatively observed coalescence times.

Seldom became this divide more apparent than during the onset of the postgenomic era, following the long-awaited arrival of genomic data. For a population-geneticist, a set of loci is a set of observed variables that can be reduced to latent variables such as population split time, geographical location or effective population size. These latent variables serve to cluster populations, which facilitated a smooth transition from single-locus to multi-locus analyses.

In contrast, for phylogeneticists, loci are traditionally the study objects, which makes redundancy analyses useful for detecting loci under selection or identifying chromosomal rearrangements, but not for phylogenetic inference. Forced back to the drawing board, phylogeneticists had to devise a novel approach for reducing the dimensionality of multi-locus data. The focus shifted from modelling among-site mutation rate heterogeneity to modelling among-site coalescence time heterogeneity (Fig. 4). A distinct discipline emerged: coalescence-based tree reconstruction (Liu et al. 2009; Jiao et al. 2021).

**Figure 4.**
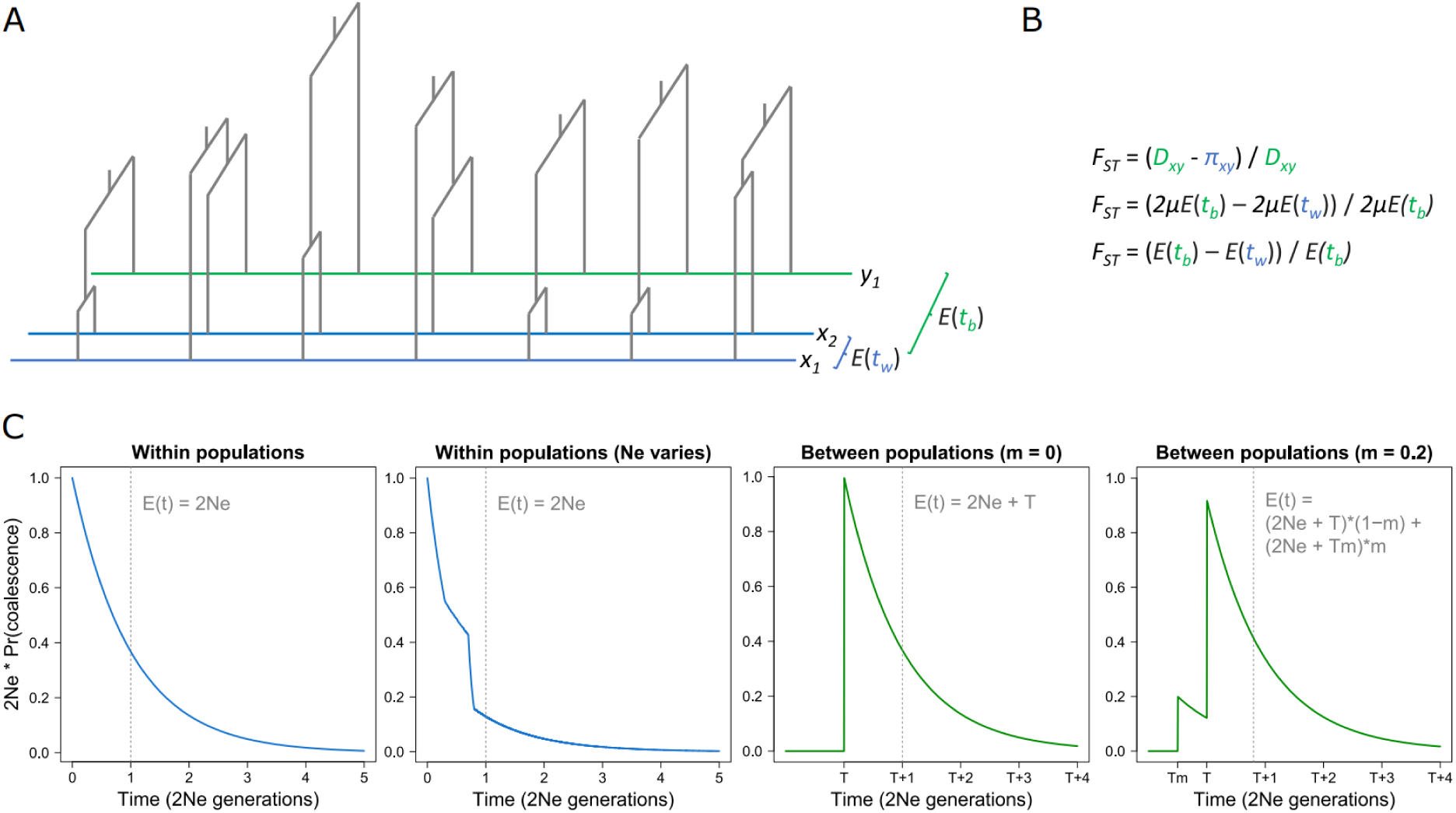
Distribution of coalescence times. **A.** Schematic diagram depicting how coalescence times, *t*, may vary among independently segregating *c-loci* along a chromosome. Population *X* is represented by two haploid individuals (*x1* and *x2*), population *Y* by a single haploid individual (*y1*). Although gene tree discordances caused by incomplete lineage sorting (ILS) are most striking (here observed for two out of seven loci), they are in fact only symptoms of the true challenge of multi-locus phylogenetic inference: how to account for variation in coalescence times across loci? This challenge remains in the absence of gene tree discordance. **B**. Assuming mutation rate constancy across lineages, *F_ST_*-estimates give the proportional difference in mean coalescence times between populations, *E(t_b_)*, versus those within populations, *E(t_w_)*. **C**. The variation in coalescence times among loci fits in principle an exponential distribution. However, population size fluctuations and gene flow events cause deviations, making the actual distribution difficult to model. For simplification, DIST only considers the expected values. The parameter m denotes the migration rate during a pulse gene flow event.

Species tree inference is a foundational goal of phylogenetics, which however only became truly attainable once single-locus stochastics could be cancelled out through multi-locus analyses (S.V. Edwards 2009). Unlike gene trees, which depict the historical relationships between non-recombining molecules, species trees depict the historical relationships between demes, be they populations or species (Maddison 1997; Liu et al. 2009). They are therefore real evolutionary phylogenies, true to the bifurcating models originally conceived by Darwin (S.V. Edwards 2009).

While a bifurcating tree accurately describes the history of molecules, with each split denoting a cell division, it may not perfectly capture the history of demes, as these do not only split but can also fuse. Fortunately, this issue can be resolved by adding gene flow edges, thereby transforming species trees into reticulate networks, also known as admixture graphs.

Even without gene flow edges, a species tree differs fundamentally from a concatenation tree, which is obtained by combining all loci into a single ‘supermatrix’ prior to tree reconstruction. Such a tree, traditionally referred to as ‘tree of individuals’ (Mountain and Cavalli-Sforza 1997; Pritchard et al. 2000), represents a descriptive model of an imaginary, average locus. Inner nodes reflect an imaginary cell division, and branch lengths are proportional to mean coalescence time, *E(t*) (Table 1, Fig. 1). A species tree, on the other hand, represents an explanatory model that can predict that imaginary average locus, as well as deviations from it (S.V. Edwards 2009). Inner nodes represent population splits, and branch lengths are proportional to population split time (*T*), often in coalescent/drift units (π = *T*/*2Ne*) (Table 1, Fig. 1).

**Table 1.** Three types of genetic trees. A species tree is an evolutionary phylogeny depicting the history of populations, and as such the ultimate goal of phylogenetic inference. In contrast, a gene tree depicts the history of molecules (i.e., haplotypes). A ‘tree of individuals’ depicts an imaginary, average gene tree. The topology of a tree of individuals is expected to be consistent with the topology of the species tree, but branch lengths denote mean coalescence times, not population split times.

|  | **species tree** | **gene tree** | **tree of individuals** |
| --- | --- | --- | --- |
| **tips** | populations/species | haplotypes | imaginary haplotypes |
| **inner nodes** | population splits | cell division | imaginary cell divisions |
| **branch lengths** | split time, $T$ | coalescence time, $t$ | mean coalescence time, $E(t)$ |

In the population-genetic paradigm, species trees inference amounts to identifying the species tree which best explains, for any population, the observed composition of alleles carried by the individuals of that population, measured in terms of allele frequencies. One aim is to explain polymorphy: why individuals which are members of the same population may yet carry different alleles (Kimura and Crow 1964). A second aim is to explain allele sharing among populations: why members of different populations may yet carry the same allele (Li and Nei 1977).

Originally performed on data sets of phenotypic traits, these analyses have a long history, featuring the development of the earliest maximum likelihood tree building methods (Cavalli-Sforza and Edwards 1967; Felsenstein 1973; Felsenstein 1973; RoyChoudhury et al. 2008; A.W.F. Edwards 2009). The underlying diffusion models strive to reconstruct allele frequency changes through time. Two recent, postgenomic additions to these models are gene flow and mutation events (Gutenkunst et al. 2009; Pickrell and Pritchard 2012; De Maio et al. 2015; Schrempf et al. 2016; Borges et al. 2022; Maier et al. 2023; Nielsen et al. 2023). Models with the latter addition do so by converting mutations models into substitution models. They are therefore known as polymorphism-aware phylogenetic models (PoMo’s), but are perhaps better understood as mutation-aware population-genetic models.

In the phylogenetic paradigm, species tree inference amounts to identifying the species tree which best explains, for any allele, the observed composition of individuals carrying that allele (i.e., population frequencies). In other words, the aim is to explain polyphyly: why individuals which carry the same allele may yet be members of different populations, and vice versa (Hudson and Coyne 2002; Rosenberg 2003; Mailund et al. 2014). Phylogeneticists traditionally do so by comparing coalescence times, through explicitly modelling the birth-death process of lineages. This is one of the aspects which sets them apart from population-geneticists, as the latter study allele frequency changes through generations without considering the fate of the lineages responsible for passing on each allele.

Owing to recombination, genealogical histories can vary greatly when sliding along a chromosome. During the transition from single-locus to multi-locus analyses, the focus of phylogenetics therefore shifted from among-site mutation rate variation to among-site coalescence time variation. This conflict becomes especially manifest when lineages coalesce further back in time than a population split. Under such conditions, known as incomplete lineage sorting or deep coalescence, gene trees arise which differ in their topology from the species tree.

Originally perceived as problematic for phylogenetic inference, challenges were quickly turned into opportunities. Multispecies coalescent (MSC) based approaches are now able to infer from the observed degree of gene tree discordance combined estimates of population split times and effective population sizes, known as coalescence units (S.V. Edwards 2009; Heled and Drummond 2010; Bryant et al. 2012; Chifman and Kubatko 2014; Liu, Xi, et al. 2015; Mirarab et al. 2021; Rivas-Gonza lez et al. 2023). This innovation transformed coalescence-based species trees from mere descriptive cladograms (Liu and Yu 2011; Mirarab et al. 2014) into explanatory models with meaningful branch lengths (Liu et al. 2010; Sayyari and Mirarab 2016). The distribution of coalescence times, often summarised using site frequency spectra, can additionally serve to extract information about migration events and population size fluctuations (Cornuet et al. 2008; Marchi et al. 2021).

It may be argued that distance-based phylogenetic methods bridge the divide between population-genetics and phylogenetics. Rather than grouping individuals for each locus by populations or by alleles, individuals are grouped pairwise, thereby side-stepping the design choice which fundamentally divides the two research fields (Fig 3). This ability to unite the two fields is evidenced by the multiple ways that a distance matrix can be constructed: expected sequence dissimilarity estimates, both within and between populations, can be calculated from allele frequencies as well as from population frequencies (Fig. S1, Table S1).

While distance-based methods are routinely used to infer gene trees and individual-level concatenation trees, they are not yet commonly applied as a tool for species tree inference. Exceptions to this rule are coalescent-based summary methods which infer species trees from matrices containing internode distances averaged over a set of gene trees (Liu and Yu 2011; Vachaspati and Warnow 2015; Liu and Warnow 2023). However, internode distances do not have a clear biological meaning, which makes it challenging to add meaningful branch lengths.

DIST infers the tree of individuals from an individual-level distance matrix containing *E(p)*-estimates, and species trees from population-level distance matrices containing either *D_xy_*, *F_ST_*-estimates or *τ*-estimates (M. J. de Jong et al. 2024). The *F_ST_*-based species tree is a descriptive model (Kitada et al. 2021). Its branch lengths indicate which proportion of lineages coalesces on average more recently than a population split (M. J. de Jong et al. 2024). This is as far as model-free distance-based approaches can lead us. The *τ*-based species tree, on the other hand, is an explanatory model, which predicts coalescence times, and thereby branch lengths of gene trees, as a function of *T* and *Ne*.

To derive *τ*-estimates from *F_ST_*-estimates, DIST makes assumptions about the underlying demographic scenario. It either assumes that the current population structure results from relatively recent population splits or instead that population size fluctuations can be ignored (M. J. de Jong et al. 2024). In the first scenario, novel mutations are negligible such that differences in coalescence time can be attributed solely to genetic drift resulting from population bottlenecks. In the second scenario, differences in coalescence times are attributed to novel mutations only, which in theory is more useful for deep divergences. Admittedly, these underlying assumptions limit the usefulness of *τ*-based species trees inferred by DIST. In contrast, *F_ST_*-based species trees are completely model-free and thus generally valid.

In the population-genetic paradigm, *F_ST_*-estimates are measures of allele frequency differences between populations. In the phylogenetic paradigm, in contrast, they quantify to what extent mean coalescence times within populations differ from those between populations (M. J. de Jong et al. 2024). Thus, consistent with being distance-based and hence mostly model-free, DIST arguably fits in either paradigm.

### Why a distance-based concatenation tree?

The key rationale behind DIST is that the individual-level concatenation tree, the tree of individuals, approximates the average gene tree, and therefore may serve to infer the species tree. A second rationale of DIST is that the concatenation tree is best inferred using a distance-based method (Sokal and Michener 1958; Fitch and Margoliash 1967; Buneman 1971; Saitou and Nei 1987; Rzhetsky and Nei 1993; Gascuel 1997; A.W.F. Edwards 2009; Pardi et al. 2010), and thus not with character-based approaches, even though the latter are in principal more sophisticated.

This design choice partly stems from practical reasons. Being based on a heuristic search algorithm, character-based approaches need a tremendous amount of computing hours to process whole-genome datasets (Kubatko and Degnan 2007; Kumar 2022; Steenwyk et al. 2023). From a more conceptual perspective, character-based approaches have been designed for analysing single, haploid loci, not for analysing multi-locus datasets of diploid individuals. This can easily lead to incorrect inferences, particularly with regard to branch lengths.

For instance, when working with relatively recently diverged populations, most observed genetic variation is expected to be ancestral, implying that population differentiation is primarily caused by genetic drift, not by novel mutations. This is in principle not a problem for phylogenetic inference, as random genetic drift can be accounted for by a lineage-specific adjustment of *E(t)* within bottlenecked populations. The risk, however, is that character-based approaches may attempt, erroneously, to fit a tree through lineage-specific adjustment of *μ* (Fig. 4). This error can be avoided by specifying a strict molecular clock, which does not allow for mutation rate heterogeneity across lineages. This forces the search algorithm to explain variation in genetic distances among individuals solely in terms of variation in *E(t)*, rather than *μ*, across lineages.

Incorrect inferences may also stem from variation across sites, rather than across lineages. When analysing a single-locus dataset, it is typically assumed that mutation rate variation across sites fits a gamma distribution (“***Γ***”) (Yang 1994). For multi-locus data, in contrast, the largest contributor to the observed variation in number of differences per site is, again, variation in *t,* not *μ* (Degnan and Rosenberg 2009; Liu, Wu, et al. 2015). In the case of population size constancy, this variation fits an exponential distribution with a mean of 2*N_e_*. Population size fluctuations cause deviations from this curve, resulting in an erratic distribution which does not fit any common probability distribution, and certainly not a gamma distribution (Thawornwattana et al. 2023) (Fig. 4).

One might therefore argue to disable mutation rate variation across sites, and to instead opt for an edge-unlinked model, in which each locus is assigned to a separate partition with unique sets of branch length (Chernomor et al. 2016). Alternatively, a computationally less demanding approach, which is less prone to the risk of overfitting, is to assume that all sites share the same mutation rates and the same coalescence times. This serves the objective of reconstructing the tree model which corresponds to the average gene tree, rather than the tree model which perfectly explains all variation in the data.

The diploid or polyploid nature of many multi-locus datasets is another potential issue which can complicate character-based tree inference. The common work-around solution is to treat heterozygous sites as ambiguous (Lischer et al. 2014), but this is dissatisfactory. The preferred solution is to randomly resolve heterozygous sites, for instance using the software vcf2phylip (Ortiz 2019). If enough loci are included in the analyses, the law of large numbers ensures that the average number of mutations between pairs of individuals is estimated accurately. The same principle is used when calculating *E(p)*-estimates between diploid individuals (see section 2).

While the suggested solutions may solve each of the forementioned issues, the safer and more scalable approach, which has been implemented in DIST, is to construct the tree of individuals using a distance-based method. Being model-free approaches, these methods can be readily applied to single-locus as well as multi-locus datasets.

## 2. FROM DATA TO DISTANCE MATRIX (STEP 1)

### Pairwise sequence dissimilarity

Genetic distances between individuals and populations can be estimated using a wide variety of measures, which often cannot be compared one-to-one, complicating useful interpretation. Even the fixation index (*F_ST_*) alone has countless definitions, with different meanings of non-extreme values (Meirmans and Hedrick 2011; Berner 2019; M. J. de Jong et al. 2024).

What is needed is a universal quantification system which ideally meets at least five criteria. The distance metric should be *i)* transferrable between different hierarchical levels (i.e., individual-level and population or species-level); *ii)* comparable across studies and organisms regardless of marker type, sequencing strategy or ploidy level; *iii*) robust to sample size variation; *iv)* a measure of a concrete concept; and *v*) comparable to predictions derived from population-genetic theory. A metric which fulfils all these requirements is pairwise sequence dissimilarity, *p*, which is the proportion of differences between two haploid sequences (Rzhetsky and Sitnikova 1996).

For diploid individuals, *p* can be approximated by calculating the mean of the four possible haplotype comparisons, which gives the expected sequence dissimilarity, here denoted as *E(p)*. Even without phase information, for a pair of diploid individuals *i* and *j*, this metric can be accurately obtained from genotype data using the formula (Box 1):

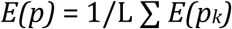

where *L* is the length of the sequence, and in which *E*(*p_k_*) is the expected distance for a single diploid site, determined using the following set of rules (Mountain and Ramakrishnan 2005; Liu et al. 2023; Xu et al. 2025) (Box 1):

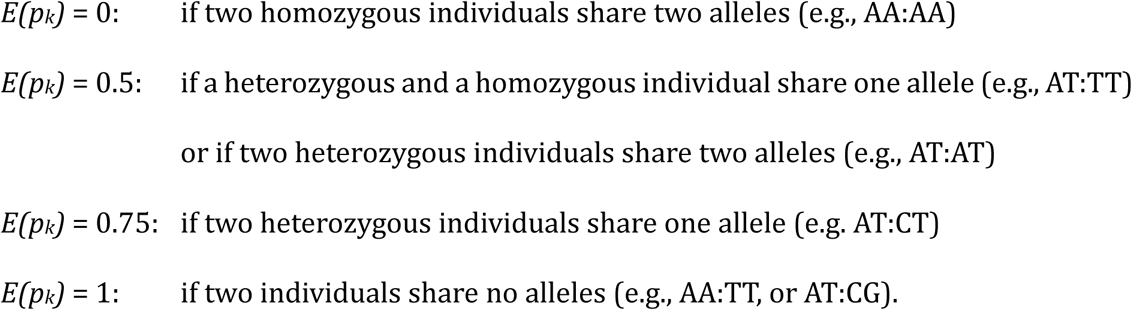

While highly correlated, *E(p)*, is not the same as a metric known as ‘allele sharing distance’ (*ASD*), which assigns to each site values of 0, 1 or 2 depending on how many alleles are shared between two diploid individuals (Mountain and Cavalli-Sforza 1997; Gao and Martin 2009). When normalised to 0, 0.5 and 1, this metric is also known as IBS, or identity by state (Chang et al. 2015; Liu et al. 2023). Discrepancies arise when two heterozygous individuals share one or two alleles, as *ASD* equals 0 and 0.5 for AT:AT and AT:CT, respectively.

A major advantage of *E(p)* is its insensitivity to ploidy levels. Since the metric gives the expected number of differences between two randomly drawn haplotypes from a pair of individuals (one from each), potential confounding effects caused by the diploid or polyploid nature of nuclear genomes are circumvented. Put differently, even though *E(p)*-estimates have been derived from diploid individuals, it is safe to assume that they represent distances between haploid individuals or haploid regions. Thus, the metric can be calculated from and compared across all regions of the genome, including autosomes, sex chromosomes and the mitochondrial genome (Fig. 5).

**Figure 5.**
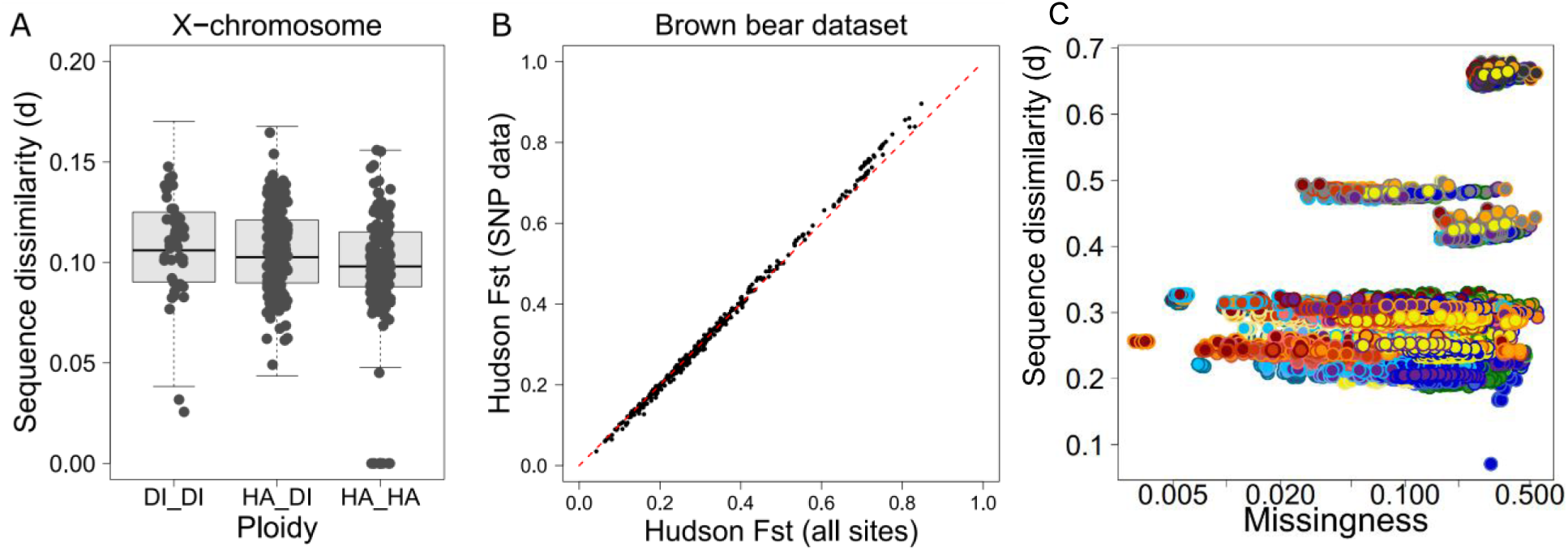
Distance calculations. **A**. DIST estimates distances between pairs of diploid individuals in terms of multi-locus expected sequence dissimilarity, *E(p)*. This metric is unaffected by ploidy, meaning it can be compared across genomic regions or individuals with different diploidy levels, as depicted here for X-chromosomal distances between males (haploid) and females (diploid). The slight difference between the categories can be likely attributed to genotype calling artifacts caused by the lower X-chromosomal depth for males. ***B***. Because the mutation rate is cancelled out when calculating relative genetic distance, *F_ST_*-estimates inferred from SNP data are expected to equal those inferred from monomorphic and polymorphic sites combined, as indeed observed for population pairs of brown bears (data from De Jong et al. 2023). Still, for comparability of *π* and *Dxy*-estimates across studies and against theoretical expectations, *E(p)*-estimates are preferably inferred from all sites, rather than from variable sites only. **C**. Ideally, no relationship should exist between missingness and sequence dissimilarity, as depicted here for a dataset of *Cervus elaphus* and closely related species (data from De Jong et al. 2025). Each datapoint represents a pair of individuals, with colour combinations indicating population pairs.

In terms of its actual meaning, *p* denotes a concrete concept, namely the observed proportion of sites which differ among two sequences. This stands in contrast to other distance metrics, in particular Euclidean distance, which estimates the distance between two individuals in multidimensional genotype space, an abstract measure with little connection to reality. One could argue that *E(p)*-estimates between diploid individuals are somewhat abstract. When focusing on a single recombination-free locus only (a coalescent locus, or ‘c-locus’), none of the dissimilarity estimates of the four possible haplotype pairs may actually agree with the mean value, *E(p)* (Box 1). However, when considering a sufficient number of unlinked loci, the law of large numbers ensures that the observed sequence dissimilarity, obtained by randomly sampling for each locus one of the four haplotype comparisons, will invariably converge to the multi-locus *E(p)*-estimate.

Concrete concepts can be translated into mathematical models. In the case of *p*, the model is a simple, linear relationship with just two explanatory variables: mutation rate (*μ*) and coalescence time (*t*). This implies that *p*-estimates for haploid loci are proportional to the coalescence time of a pair of sequences. In the case of diploid data, single-locus *E(p)*-estimates are proportional to the expected coalescence time, *E(t)*, averaged over the four haplotype comparisons. Multi-locus *E(p)*-estimates are proportional to *E(t)* averaged over loci. For high *E(t)*-values, when multiple mutations per site cause *E(p)* to be an underestimate, a correction method can be applied to restore the linear relationship between *E(p)* and *E(t)* (Kimura 1980).

No such straightforward relationship exists between *t* and other distance metrics. As a numerical example, imagine a small dataset of two loci and two diploid individuals. If inbreeding converts the allele frequency differences between the two individuals from 0.5 and 0.5 to 0 and 1, the multi-locus *E(p)*-estimate remains constant, namely 0.5, while the Euclidean distance increases from 0.71 to 1. This explains why in a tree of individuals inferred from a Euclidean distance matrix, inbred individuals typically have elongated branches and thus falsely appear as outgroup individuals with deep coalescent times.

Because *E(p)* denotes a concrete concept with a real biological meaning, it is no surprise that the metric allows to unite all major population-genetic parameters. Simply by averaging over pairs of individuals, sequence dissimilarity can be directly translated into population-level diversity and divergence estimates, namely nucleotide diversity (*π*) and absolute genetic distance (*D_xy_*). As discussed in more detail further below, a comparison between these parameters gives net divergence (*D_a_*, or Nei’s *D*) and relative pairwise population divergence (*F_ST_*). This allows to combine all these standard population-genetic measures into one unifying framework, and thereby also to display them in a single dendrogram.

Conveniently, all these estimates can be derived irrespective of sample size variation across populations. Because nuclear genomes are comprised of huge numbers of c-loci (Doyle 2022), accurate estimates of *π*, *D_xy_*, *D_a_* and *F_ST_* can, in theory, even be obtained with sample sizes as low as a single diploid individual per population, provided panmixia. However, in practice, it is still advisable to obtain data for at least a few individuals per population. This allows to validate the panmixia-assumption, which is frequently violated by fine-scale population structure and/or inbreeding (i.e., close-kin matings).

### Deriving *He, π*, *F_ROH_*, *D_xy_*, *D_a_* and *F_ST_* from *E(p)*

When averaging *E(p)* over all pairs of individuals within a population, we obtain its mean nucleotide diversity, *π_xy_* (Nei and Li 1979). When averaging *E(p)* over all pairs of individuals between two populations, we obtain *D_xy_* (Nei 1987: 10.20):

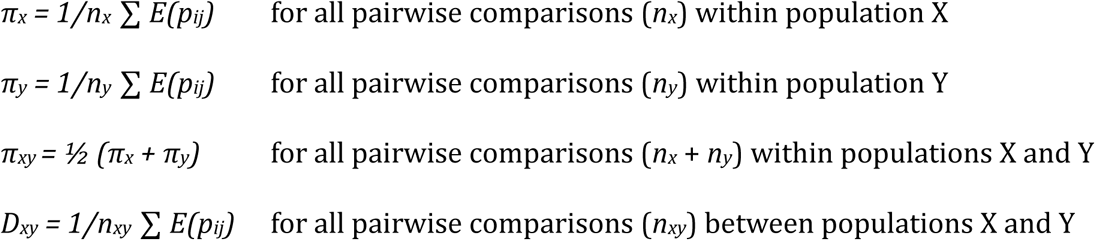

The *π_x_-* and *π_y_-*estimates give the mean observed distance within a population (Box 2). For panmictic populations, this nucleotide diversity is expected to equal the genome-wide heterozygosity, defined as the proportion of heterozygous sites within a genome, of each individual (i.e., *π* = *He*). Deviations are indicative of inbreeding caused by close-kin mating (*F_IS_*), which hence can be calculated as: *F_IS_x_*: (*π_x_* – *He*)/*π_x_*. *F_IS_* is expected to be equivalent to the genome-wide proportion of runs of homozygosity, *F_ROH_*. The average of *π_x_* and *π_y_* is known as *π_xy_* (‘pixy’).

Whereas *D_xy_* denotes the absolute distance between populations, the relative distance can be estimated using the fixation index, *F_ST_*. This fixation index can be obtained using the definition of Hudson et al (1992), which is insensitive to sample size variation (Bhatia et al. 2013; M. J. de Jong et al. 2024). This Hudson *F_ST_* metric normalises net divergence (*D_a_*), which is the difference between *D_xy_* and *π_xy_* (Takahata and Nei 1985), and which is approximated by Nei’s D (M. J. de Jong et al. 2024):

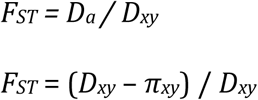

### Why distance calculations should include monomorphic sites

Population-geneticists often perform analyses on data sets which contain polymorphic sites only, i.e., SNP datasets. (Leache et al. 2015; Schmidt-Lebuhn et al. 2017). The rationale is that monomorphic, or non-variable, sites do not affect the relative distances between individuals within the dataset and hence can just as well be omitted. For example, if an unbiased selection of polymorphic sites indicates that the genetic distance of individual *i* to individuals *j* and *k* equals 0.1 and 0.2 respectively, and if the proportion of polymorphic sites is 0.01, then based on all sites the genetic distances of individual *i* to individuals *j* and *k* equals 0.001 and 0.002, respectively. Irrespective of including or excluding monomorphic sites, the distance *p_ij_* is half the distance *p_ik_*. This means that a tree of individuals generated from a SNP-based distance matrix will be identical to a tree of individuals generated from a distance matrix inferred from all sites (i.e., monomorphic and polymorphic), except for the scale of the branch lengths (Fig. 5). It also implies that accurate *F_ST_*-estimates, and thereby species trees, can be obtained from SNP data alone (see section 3).

Still, there are two important reasons why *E(p)*-estimates are ideally calculated over all sites, not from SNP datasets. First, SNP datasets seldom contain a fully unbiased selection of variable sites. This is especially the case when data has been produced using SNP beadchips, which typically target a non-random selection of variable sites with high allele frequencies in reference populations (Leache and Oaks 2017). Ascertainment bias is less of a concern when extracting SNPs from genome-wide resequencing data or random reduced representation libraries (e.g., RADseq data), but biases may still involuntarily be introduced during SNP filtering.

One potential source of bias is a filter on minor allele frequency (MAF). Imagine, for instance, using a MAF-filter of 5% for a dataset of hundred individuals of which two belong to a distant outgroup. This will remove all SNPs which are private to the outgroup, resulting in an underestimate of the genetic distance of the outgroup relative to the ingroup, causing the output of phylogenetic or structure analyses to be unreliable. As a second example, a LD-filter favours the selection of uncorrelated SNPs, and thereby may magnify discordant signals resulting from incomplete lineage sorting or gene flow. A third commonly applied filter is the number of alleles per site. SNP datasets usually contain biallelic SNPs only, not multi-allelic sites with three or four alleles per site. Removing multi-allelic sites may affect genetic distances between individuals unequally (Bertels et al. 2014).

The second reason why genetic distances should not be calculated from variable sites only, is that the omission of monomorphic sites compromises how insightful the absolute distance estimates are. When calculated from variable sites only, absolute distance estimates within (*He*, *π*) and between (*D_xy_*, *D_a_*) populations cannot be compared across studies, nor to expectations from population-genetic theory.

The inclusion of monomorphic sites when calculating *E(p)*-estimates does not necessarily mean that the full data set needs to be considered. The law of large numbers ensures that a random, unbiased selection of sites also suffices to obtain an accurate and precise estimate of genomic-wide genetic distance. Such a random selection of sites can be obtained by randomly thinning the data, for instance by using the ‘thin’ option of vcftools (Danecek et al. 2011). As a rough rule of thumb, distance estimates can be reliably obtained from a random selection of sites containing 1% of a 2Gb genome (Fig. S2), which implies that DIST can also be applied to RADseq data.

## 3. FROM DISTANCE MATRIX TO TREE OF INDIVIDUALS (STEP 1)

### Evaluating a tree of individuals

A individual-level distance matrix typically serves to infer a tree of individuals, using a hierarchical clustering method such as the bioNJ or OLS algorithms (Gascuel 1997; Gascuel et al. 2001; Desper and Gascuel 2004). In the DIST workflow, however, the tree of individuals is just an intermediate output, assumed to approximate the imaginary average gene tree. Users are, nevertheless, encouraged to observe the tree of individuals, as it allows to validate the underlying distance matrix, and to read the population-genetic parameters used by DIST for inferring the species tree.

The tree of individuals should not be taken at face value but instead needs to be validated first by evaluating the fit between the model (the tree) and the underlying data (the genotypes). Being distance-based, this validation process consists of answering two separate questions (Pardi and Gascuel 2012): 1.) does the distance matrix accurately represent the true distances between individuals, and 2.) does the tree accurately represent the distance matrix?

A straightforward approach to answer the second question is to compare the tip-to-tip path lengths in the dendrogram (*l*) to their corresponding entries in the underlying distance matrix, *E(p)* (Buneman 1971; Malinsky et al. 2018; de Jong et al. 2023). We can define the residual error (Ɛ), or ‘stress’, of a dendrogram as the proportional distance between *l* and *p*:

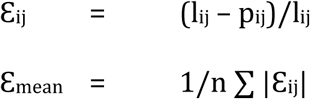

A positive *Ɛ_ij_*-value indicates that the genetic distance between two tips, *i* and *j*, is smaller than suggested by the path lengths of the dendrogram. A negative *Ɛ_ij_*-value, in contrast, indicates that the genetic distance between *i* and *j* is larger than suggested by the dendrogram. The mean over all absolute values combined, *Ɛ_mean_*, is a measure for the overall fit between the model (the bifurcating tree) and the data (the distance matrix), with zero denoting a perfect fit. A heatmap depicting the residual error for each pair of individuals should ideally be presented alongside the dendrogram, as this allows to evaluate the fit between the tree of individuals and the underlying distance matrix (Cavalli-Sforza and Piazza 1975).

Residual error is essentially a measure of non-additivity (A.W.F. Edwards 2009), which may arise from two main sources: gene flow events, or artefacts introduced by sequencing and/or genotype calling errors. Residual error cannot be attributed to gene tree discordances resulting from deep coalescence (i.e., incomplete lineage sorting), nor to mutation rate variation across lineages, as both factors do not affect the additivity of *E(p)*-estimates. Thus, in the absence of gene flow, any observed residual error either reflects sequencing or genotyping artefacts, or inconsistency of the clustering algorithm.

If, instead, gene flow did occur, the distance matrix is not additive, in which case no bifurcating tree can be reconstructed which perfectly fits the data (see also section 5) (Cavalli-Sforza and Piazza 1975; Peter 2016). Residual error is to be expected, irrespective of the consistency of the clustering algorithm, and more generally, irrespective of the phylogenetic approach used to construct a bifurcating tree (i.e., whether distance-based, character-based or coalescent-based).

Nevertheless, certain clustering algorithms perform better than others, and large variation may exist in the residual errors produced by different clustering algorithms (de Jong et al. 2023). The lowest residual errors are expected when using the OLS algorithm, as this algorithm has been particularly designed to minimise residual error, while other methods aim to optimise other parameters (e.g., BME minimising the sum of all branch lengths) (Rzhetsky and Nei 1993; Gascuel et al. 2001; A.W.F. Edwards 2009; Pardi et al. 2010; Vaz et al. 2021).

### Using the tree of individuals to validate the distance matrix

The residual error of a dendrogram allows to validate the dendrogram, but how to validate the distance matrix? In other words, how to know if the genotypes (or genotype likelihoods) have been called reliably and subsequently distances calculated correctly? Apart from building in checkpoints during the genotype calling process (Fig. 5), we can use for this purpose the criterium of ultrametricity.

When distance calculations are performed for a large and unbiased set of loci, and assuming absence of gene flow as well as absence of mutation rate heterogeneity, multi-locus distances are not only additive but even ultrametric. This translates to equal root-to-tip distances, which creates perfectly circular structures when the tree of individuals is displayed in the unrooted format.

The assumption of mutation rate constancy is frequently violated for single-locus data, either due to selective pressures (i.e., in case of gene or protein sequences) or due to the small size of the sequences, which lead to stochastic variations. In contrast, for distance matrices constructed for individuals from relatively closely related species, mutation rate heterogeneity is much less of a concern. Local mutation rate variations are cancelled out when analysing numerous loci randomly distributed across the genome. We therefore are left with global, genome-wide mutation rates, which vary little across closely related lineages (Zhang et al. 2023), which justifies the use of distance-based approaches (Rzhetsky and Sitnikova 1996).

For relatively recent population splits, which occurred during the Holocene or even Late Pleistocene, between-population distances (*D_xy_*) often barely exceed within-population distances (*π*). As a consequence, population differentiation can often only be reliably detected if distance estimates are precise and accurate (Bowcock et al. 1994). Validation of the distance matrix using the ultrametricity criterium therefore is crucial for ensuring that true phylogenetic signals are not lost in the noise introduced by data artifacts. One potential complication is that deviations in root-to-tip distances may not necessarily be artifacts but may also be indicative of gene flow (see section 5). However, gene flow cannot explain large deviations between individuals from within panmictic populations.

Potential causes for data artifacts are systematic sequencing error, genotype calling bias, SNP ascertain bias (e.g., MAF-filter and LD-pruning), usage of a distance metric other than *d*, or insufficient data. In the case of the latter, the number of loci from which genetic distances have been estimated does not suffice to fully cancel out single-locus stochastics.

The variation in *E(p)*-estimates, both within and among populations, can be quantified by calculating standard deviations. A t-test can serve to determine whether *E(p)*-estimates between populations A and B differ significantly from those between populations A and C. This test may however only be applied if each individual is an independent replicate of its panmictic population. This is the case if individuals are unrelated, and if genotypes have been called, or genotype likelihoods inferred, for each individual independent from other individuals. This means that if using the software ‘bcftools call’ for genotype calling, each individual should be assigned to its own, unique group using the ‘group-samples’ option.

If standard deviations are low, the individual-level distance matrix may be further reduced to a population-level distance matrix, containing *D_xy_* and *π*-estimates. Such a population-level distance matrix contains mean absolute distances between population-level consensus sequences, can be used to infer a ‘tree of populations’. This type of trees should however not be mistaken for a species tree, as inner nodes still represent an imaginary cell division, and as branch lengths are still proportional to *E(t)*, not *T*.

### Interpreting a tree of individuals

A tree of individuals constructed from pairwise *E(p)*-estimates is an all-in-one summary of genomic data, depicting many population-genetic parameters at once. This includes nucleotide diversity (*π*), genome-wide heterozygosity (*He*), absolute genetic distance (*D_xy_*) and relative genetic distance (*F_ST_* and *f^2^*) (Fig. 6) (M. J. de Jong et al. 2024).

**Figure 6.**
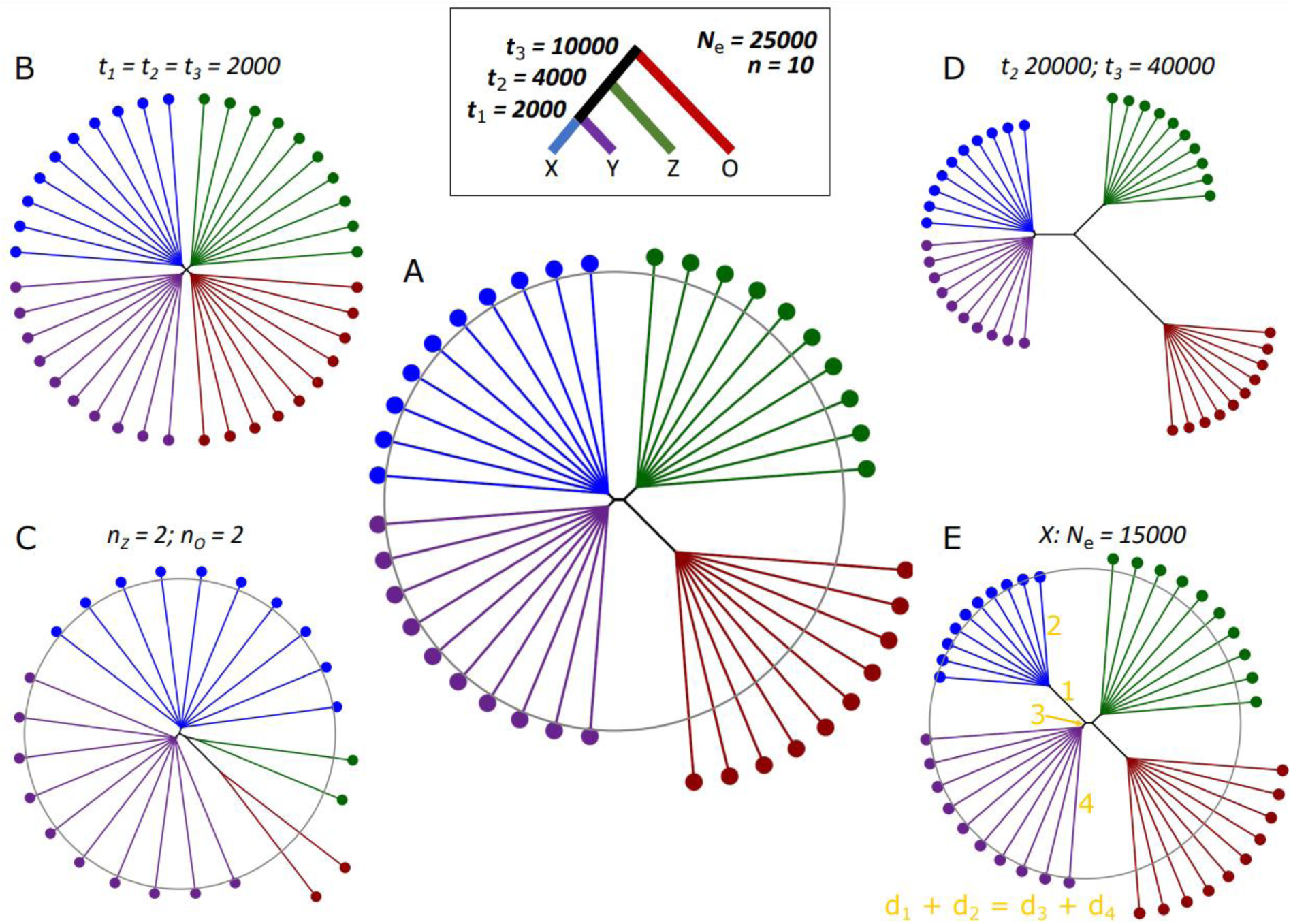
*Concatenation trees for various demographic scenarios*. **A**. BioNJ-dendrograms, inferred from a simulated distance matrix with multi-locus *E(p)*-estimates. The distance matrix has been predicted for the demographic scenario indicated in the inset, assuming a mutation rate of 10^−8^ mutations per site per generation. The circle has been added for visual interpretation, and serves as isoline. Note that individuals within panmictic populations are expected to be equidistant to each other, which provides a simple criterium to evaluate the accuracy of a concatenation tree. ***B***. Idem as A, but with a hard polytomy: *t_1_* = *t_2_* = *t_3_* = 2000 generations. ***C***. Idem as A, but with smaller sample sizes for populations *Z* and O. Note that due to the unbalanced sample sizes, which create obtuse and astute angles between branches, population Z can be mistaken to cluster with O. ***D***. Idem as A, but with *t_2_* = 20000 and *t_4_* = 40000. ***E***. Idem as A, but with *N_e_* = 15000 for bottlenecked population X. The population bottleneck reduced mean coalescence time within population X, making these individuals more similar to each other (small *d_1_*), without affecting their distance to individuals of other populations (*d_1_* + *d_2_* = *d_3_* + *d_4_*). This insensitivity of *Dxy* to population size fluctuations is a useful property, particularly for split time inference.

When using *E(p)* as distance metric, and assuming the inclusion of sufficient loci to cancel out single-locus stochastics, the shape of an unrooted tree of individuals is predictable. In the absence of gene flow, and assuming panmixia, tips of unrooted individual-based dendrograms will cluster into neat circles with radii corresponding to ½*π* (Fig. 6). This split is actually multifurcating, as within panmictic populations equal E(t)-estimates are expected for all pairs of individuals, which is approximated by a series of bifurcating splits which with inner branch lengths converged to zero.

For large populations which split relatively recently, between-population differences barely exceed within-population differences (i.e., *π ≈ D_xy_*), such that unrooted dendrograms will assume an almost perfect circular shape, with a radius *r* = ½*D_xy_* (Fig. 6). With increasing split time, this circular shape will decompose into diverging semi-circles, with each semi-circle corresponding to a distinct, panmictic population.

If the dataset contains bottlenecked populations, these individuals will be conspicuous owing to their relatively short terminal branches, which indicate low genetic variation within the population (low *π*, high *F_ST_*). These short terminal branches are compensated for by a longer inner branch, of which the length will be proportional to the amount of genetic variation lost after the population split (Fig. 6).

While bottlenecked populations clearly stand out due to their short external branches, the true outgroup populations may be hardly noticeable, as even after considerable split times *D_xy_* may barely exceed *π_xy_* (Bowcock et al. 1994). This may at first appear puzzling. How can genetic distances within populations be barely shorter than genetic distances between populations?

The key to understanding is that path lengths are proportional to mean coalescence time, *E(t)*, not to population split time, *T* (Table. 1). The mean coalescence time of two haplotypes sampled within a diploid population is 2*N_e_* generations, which even for moderately sized population corresponds to thousands of generations (Hancock and Blackmon 2020). For deep population splits (e.g., dating to the Pliocene or older era), novel mutations greatly outweigh the coalescence probability in the ancestral populations (2*Ne_anc_*). But when studying more recent processes, such as Holocene or Late Pleistocene populations dynamics for species with *Ne* > 10000 and *G* > 5 (*G* for generation time in years), it is the other way around: 2*Ne_anc_* greatly outweighs *T*. For these study systems, the observed genetic variation mainly reflects ancestral variation, with relatively few novel mutations having accumulated after a population split (Bowcock et al. 1994).

The implication is that for study systems of medium to large populations (high *Ne*), even the smallest inner branches may be indicative of considerable periods of isolation. This is an important difference compared to single-locus trees, for which short inner branches are usually associated with low support values, indicating that alternative topologies cannot be ruled out. For multi-locus trees, in contrast, internal branches can be short yet robust.

Genetic clusters separated by short internal branches are arguably better observed when the tree is depicted using the unrooted format rather than the phylogram format. However, a disadvantage of the former is that outgroups are more difficult to recognise. When sample sizes are unbalanced, the angle between the outgroup and the underrepresented ingroup becomes acute, and the angle between the outgroup and the overrepresented ingroup obtuse. This may give the false impression that the underrepresented ingroup clusters with the outgroup. To aid visual interpretation and to make outgroups more conspicuous, it is often useful to add one or multiple concentric circles which can serve as ‘isolines’ (Fig. 6) (de Jong et al. 2025).

In the presence of recent or ongoing gene flow, such that admixed alleles did not have sufficient time yet to distribute homogenously throughout the recipient population, the genetic distances between pairs of individuals of recipient and donor populations will vary extensively. As a result, individuals of admixed populations will form in the dendrogram, between the two donor populations, a fan-like collection of separate branches, rather than a distinct unit.

A homogenously admixed population, which received gene flow during a more ancient admixture event, can be recognised from deviations in branch lengths. If this population introgressed from a distant outgroup a relatively small proportion of genetic material, it will have (slightly) enlarged branch lengths relative to its sister populations. When proportions of introgressed DNA are higher, the admixed population will cluster as an outgroup relative to its sister populations. In all scenarios, the residual error heatmap will indicate that for pairwise population comparisons involving the admixed population, the path lengths of the dendrogram are not consistent with the true underlying values in the distance matrix. Therefore, calculating residual error can serve as an indirect test of gene flow (Cavalli-Sforza and Piazza 1975; Peter 2016; de Jong et al. 2023).

## 4. FROM DISTANCE MATRIX TO SPECIES TREE (STEP 2)

### Reconstructing the species tree

The second step of the DIST workflow is to infer the species tree from the distance matrix, using the following algorithm:

1. Summarize *E(p)*-estimates into mean values within and among populations: *π, D_xy_*.
2. Infer relative genetic distances between population pairs using the formula of Hudson (1992): *F_ST_* = (*D_xy_* – *π*_xy_)/*D_xy_*, with *π*_xy_ being the mean of *π*_x_ and *π*_y_.
3. Infer estimates of coalescent units between pairs of populations, *τ_xy_*, using either, in the case of recent population splits, the formula *τ_xy_* = -ln(1 – *F_ST_*), or alternatively, in the case of population size constancy, the formula *τ_xy_* = *F_ST_* / (1 – *F_ST_*).
4. Reconstruct the species tree from the distance matrix with *τ_xy_*-estimates, using a clustering algorithm, such as bioNJ or OLS.
5. Estimate the degree of gene tree discordance using the formula: *P_discordance_* = ⅔· (1 – *F_ST_*).

This section explains the population-genetic theory underlying this algorithm.

## Theoretical predictions for *π* and *D_xy_*

Multi-locus *E(p)*-estimates quantify the proportion of sites for which two individuals carry a different allele. Assuming sufficient data to cancel out single-locus stochastics, and assuming global mutation rate constancy across lineages, these proportions are given by the expected values of the probability distributions of coalescent times (Fig. 4). The species tree can be inferred through establishing which combinations of *Ne* and *T* can explain these mean coalescent times.

To solve this inverse problem, we only need to consider a few basic population-genetic formulae. In case of not-too-deep coalescence times, for which *t* < 1/μ generations (Slatkin 1993), the expected sequence dissimilarity between any two haploid sequences for any locus *k*, is given by a simple linear equation:

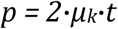

In here, *µ_k_* denotes the locus-specific neutral mutation rate per site, and *t* denotes time to coalescence (Hancock and Blackmon 2020). For long evolutionary time scales, when the linear relationship between sequence dissimilarity and coalescence time breaks down owing to parallel and back mutations, the linear relationship can be restored using a correction method, such as the Kimura-Two-Parameter correction method (Kimura 1980).

This linear expression, which lays the foundation of all formulae presented below, is very intuitive. It indicates that the difference between any pair of sequences depends on the number of mutations which both lineages – hence we multiply by 2 – accumulated after their split. This number of mutations is simply given by the number of time units since the population split multiplied by the expected number of mutations per time unit. The time unit can either be year or generation, but for comparability between species, with potentially vastly different generation lengths, it is more useful to think in terms of number of generations.

For multi-locus data, the explanatory and dependent variables are summary estimates averaged over unlinked loci, such that the expected sequence dissimilarity is a function of the global, genome-wide mutation rate, *μ*, and the mean coalescence time, *E(t)*:

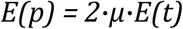

With this single and simple formula, we can predict genetic distances within (*π*) and between (*D_xy_*) populations (M. J. de Jong et al. 2024). Assuming constancy of global mutation rates across closely related lineages, we only need to consider how E(t) is affected by population size (*Ne*) and by population split time (*T*).

Coalescent theory predicts that the mean time to coalescence within populations, *E(t_w_)*, equals the effective population size, which is 2*N_e_* for diploid populations. This is for the same reason as why the expected number of dice rolls before throwing a predefined value depends on the number of sides that the dice contains. For instance, in case of an ordinary 6-sided dice, it takes on average six throws before the dice ends up on the desired side.

Having determined the mean time to coalescence within populations, *E(t_w_)*, we can infer the expected sequence dissimilarity within populations (*π*):

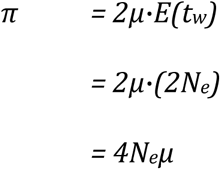

When, instead, comparing sequences drawn from two different populations, the expected level of sequence divergence (*D_xy_*) is the sum of two factors. First, the nucleotide diversity of the ancestral population (*π_anc_*) defines the base level of *D_xy_* at the time of the population split. Second, *D_xy_* increases through mutations accumulated after the population split, *T* (Takahata and Nei 1985; Wakeley 2000; Nachman and Payseur 2012; Crawford et al. 2015; Vijay et al. 2017; Delmore et al. 2018; Hancock and Blackmon 2020; M. J. de Jong et al. 2024). Note that T should not be confused with coalescence time, *t*. We thus obtain:

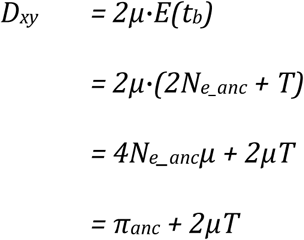

Because mutation rates in nuclear genomes are low, the factor *2μT* has a negligible effect on *D_xy_* when population splits occurred relatively recently. For instance, assuming *μ* = 10^−8^ and *π_anc_* = 0.002, it takes up to 10^4^ generations for *D_xy_* to increase from 0.002 to 0.0021 only. This implies that for populations which split during the Holocene or Late Pleistocene, and which have a generation length of several years, the factor 2*μT* can often safely be ignored, such that the formula simplifies to:

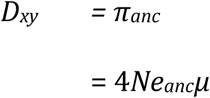

This indicates that *D_xy_* depends only on the effective population size of the ancestral population, *Ne_anc_*, and not on the effective population size of extant populations. The reason is that loss of variation due to random genetic drift makes individuals within a population more similar to each other but does not make these individuals more different to individuals of other populations. In other words, random genetic drift affects sequence dissimilarity within populations, such that *π* decreases, but not sequence dissimilarity between populations, such that *D_xy_* remains constant. Thus, unlike *F_ST_*-estimates, multi-locus *D_xy_*-estimates are not affected by random allele frequency fluctuations following a population split (Cavalli-Sforza and Piazza 1975; M. J. de Jong et al. 2024).

The conclusion that multi-locus *D_xy_*-estimates are not affected by random genetic drift also pertains to a process known as ‘genetic draft’: allele frequency changes caused by linked selection. Linked selection, whether positive (‘selective sweep’) or purifying (‘background selection’), causes a reduction of genetic diversity in the genomic region surrounding the locus under selection, but on average does not alter the genetic distance (i.e., *D_xy_*-estimate) relative to other populations (Hudson et al. 1987; Birky and Walsh 1988; Menno J. de Jong et al. 2024). This means that, conveniently, *<u>D</u>_xy_*-estimates can be safely inferred from genomic datasets, even if most regions in the genome are affected by linked selection (Harris 2018; Pouyet et al. 2018).

### Theoretical predictions for *F_ST_* and *τ*

From the perspective of coalescent theory, genetic drift decreases the coalescence time (*t*) between haplotypes samples from within the same population. In other words, genetic drift is accounted for in the formula *p = 2μt* through adjustment of *t*. A pair of haplotypes drawn from a small population, which lost much of its genetic diversity as a result of genetic drift, are expected to have a lower *t* than a pair of haplotypes drawn from a larger population (Nachman and Payseur 2012; Delmore et al. 2018; Lucena-Perez et al. 2021).

The reduction of *π* due to the genetic drift, also known as the Hagedoorn effect (Hagedoorn and Hagedoorn, A. C. 1921), depends on two factors: effective population size (*Ne*) and population split time (*T*). If assuming absence of novel mutations, and non-overlapping generations (Wright-Fisher model), nucleotide diversity can therefore be expressed as a function of *π_anc_*, *N_e_* and *T*:

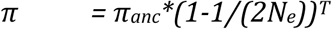

In here, *T* represents the time since the population split, assuming complete isolation of demes after their isolation. In the case of subsequent gene flow events, it may be replaced with *T_e_*, which denotes effective separation time, analogous to effective population size (Mountain and Cavalli-Sforza 1997).

Since it is usually impossible to attribute the loss of genetic variation to an exact combination of *N_e_* and *T*, the best we can do is to express population split times in multiples of their haploid effective population size, also known as coalescent units (*τ*):

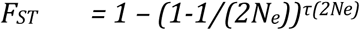

Inverting the above formula, we can infer *τ* from *F_ST_*, as follows (M. J. de Jong et al. 2024):

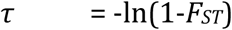

It should be cautioned that this formula provides a good approximation only in case of recent population splits, when novel mutations are negligible. For deep population splits, when divergence can be attributed to novel mutations mainly such that population size changes play a negligible role, the relationship between *τ* and *F_ST_* is approximated by (M. J. de Jong et al. 2024):

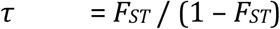

In either scenario, *F_ST_* represents the inverse of deep coalescence, namely the proportion of haplotypes that coalesce more recent than the population split (M. J. de Jong et al. 2024). It is worth noting that both types of proportions are not determined by how fast haplotypes are evolving, but instead only depend on the factors *T* and *Ne*. This implies that *F_ST_*-estimates, and thus coalescent units, are independent of the mutation rate:

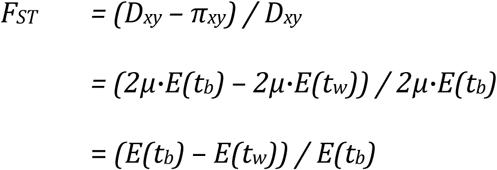

Consequently, *F_ST_*- and *τ*-estimates can, in principle, be inferred from SNP data sets, from which monomorphic sites have been excluded, provided absence of filter settings which introduce involuntarily biases (see section 2).

### Quantifying gene tree discordance

Reducing all genome-wide variation into a distance matrix with point estimates does not mean that the conflict in the data is lost. The species tree can predict the extent the degree of gene tree discordance resulting from incomplete lineage sorting.

Coalescent theory predicts that two-thirds of deep coalescing haplotype pairs are expected to be discordant (Jiao et al. 2021). Since *F_ST_* represents the inverse of deep coalescence, we can use *F_ST_* to infer the degree of gene tree discordance within genomes (M. J. de Jong et al. 2024):

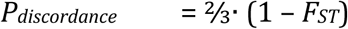

This formula assumes that gene tree discordance is caused by incomplete lineage sorting only. In the next and last section, we will discuss how distance matrices allow to determine if gene tree discordance is caused, instead, by gene flow events, and in which case a bifurcating species tree is a contrived model.

## 5. FROM SPECIES TREE TO RETICULATE NETWORK (STEP 3)

### From species tree to reticulate network

In the absence of gene flow, a bifurcating species tree can perfectly capture the relationships between demes, thereby accounting for gene tree discordance caused by incomplete lineage sorting. In contrast, in the presence of gene flow, no bifurcating tree exists which provides a complete and accurate picture of the relationships between demes (Malinsky et al. 2018). Fortunately, the fit between the data and the model can be improved by adding gene flow edges. This optional third step of the DIST workflow transforms species trees into reticulate networks, which unlike bifurcating trees can contain multiple incoming edges per node (Makarenkov and Legendre 2004; Velasco and Sober 2010).

Population-geneticists refer to reticulate networks, not be confused with a splits network (Huson and Bryant 2006), as admixture graphs (Peter 2016; Peter 2022; Maier et al. 2023; Nielsen et al. 2023). They use the term ‘admixture edge’ for both incoming edges rather than differentiating between a gene flow edge and a main edge. This is indeed arguably more correct, as the exact meaning of two edges depends on the demographic context, particularly on the magnitude and duration of migration.

Following a gene flow event, introgressed alleles will randomly replace existing alleles of the recipient population and thereby effectively overwrite genetic information (Koonin 2009). This will alter the proportion of concordant and discordant loci. For any quartet of haplotypes, there are three unbalanced quartet topologies, with the concordant topology *q1* being the most frequent, and with discordant topologies *q2* and *q3* occurring in equal frequencies (i.e., *q1* > *q2* = *q3*). Gene flow events will alter the relative frequencies of the three topologies and cause an imbalance of the frequencies of *q2* and *q3*, which can be detected using gene flow tests such as the *D*-statistic.

In the case of a strong or persistent gene flow event, one of the two alternative topologies, *q2* or *q3*, might become more frequent than the topology *q1*. Following that transition, the inferred species tree will correspond to the new population structure which came into existence with the onset of gene flow. The gene flow edge now represents the ancestral population structure, prior to the removal of this barrier.

### Testing for admixture using the *d3*-score

Gene flow events can be inferred by testing for deviations from equidistance of an outgroup to two or more ingroup lineages. While the Four Taxon Test (i.e., *D*-statistic) is more popular, gene flow can also be inferred from a comparison between three lineages.

Without gene flow, and assuming mutation rate uniformity across lineages, two ingroup populations, *A* and *B*, are expected to be equidistant to any outgroup population *C* (Mualim et al. 2021) (Fig. 7). In the case of gene flow between an outgroup population and one of the ingroup populations (e.g., from *B* to *C* and/or vice versa), this equidistance disappears. The expected distances for any population triplet can thus be summarised as follows:

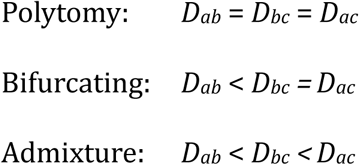

**Figure 7.**
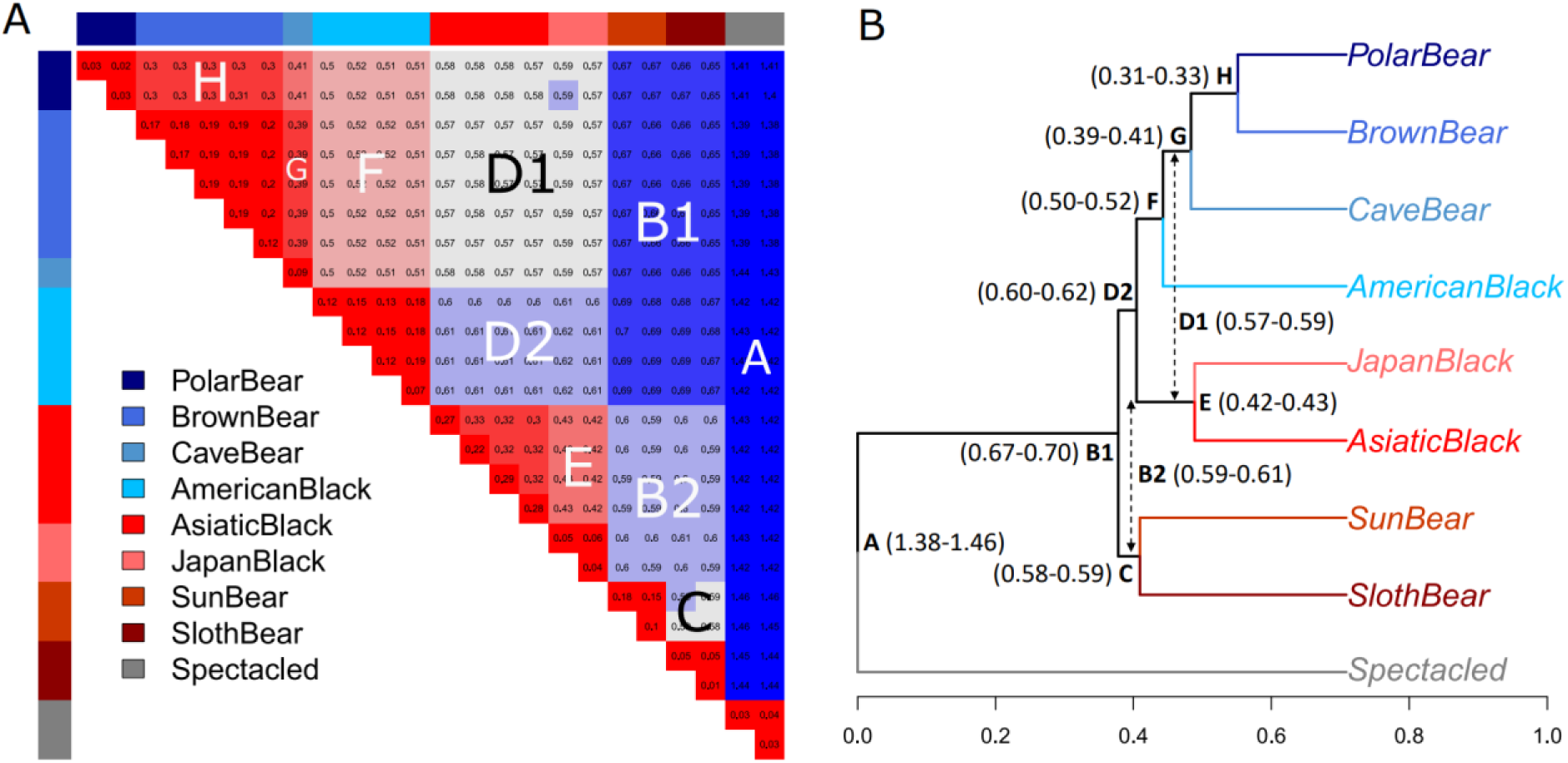
Non-additivity indicates gene flow events among bear species. DIST infers gene flow events from multi-locus distance matrices by testing for deviations from equidistance of an outgroup lineage to two or more ingroup lineages. **A**. This figure is a remake of figure 5 in the classic paper of Cavalli-Sforza and Piazza (1975), derived from real-world data of bear species. The matrix depicts genome-wide sequence dissimilarity, *E(p)*, in percentages, between bear species, dissected into blocks of equal distances. Blocks B2 and D2 deviate from the pattern expected for strictly bifurcating trees. **B**. A species tree depicting absolute genetic distances (*D_xy_*) between bear species, with node labels corresponding to the blocks in the distance matrix. Arrows indicate violations of additivity, suggestive of gene flow

Based on these expectations, and using a formula similar to the *D*-statistic, Hahn and Hibbins (2019) suggested the following three-sample test to detect admixture:

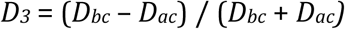

An alternative method, which does not require as input the topology of a bifurcating tree, is to calculate for each population triplet a measure that we will refer to as the ‘*d_3_*-score’ (lower case ‘d’):

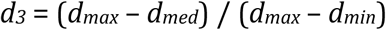

In here, *d_max_*, *d_med_*, and *d_min_* denote the maximum, median, and minimum of the three *D_xy_*-values for a given population triplet, which for above example correspond to *D_ac_, D_bc_* and *D_ab_*, respectively. The *d_3_*-statistic can thus range between 0 and 1, with 0 indicating no deviation from equidistance (i.e., *d*_max_ = *d*_med_) and 1 indicating that the intermediate population is equally distant to the other two populations. Assuming mutation rate uniformity across lineages, and absence of subsequent gene flow events, the *d_3_*-score remains constant through time (Box 3). If multiple individuals have been sampled per population, a *t*-test allows to evaluate whether the difference between *d_max_* and *d_med_* is significant. Based on the parsimony principle, by assuming that a scenario entailing few gene flow events is more likely, the consistency of the *d_3_*-score across population triplets allows to infer the timing of the gene flow event relative to population splits (Malinsky et al. 2021).

The *d_3_*-score does not contain information about the direction or magnitude of gene flow. Under certain conditions, admixture proportions can be quantified with a measure that we refer to as the *d_4_*-score.

### Quantifying admixture using the *d_4_*-score

It is a central theorem of population-genetic theory that the substitution rate is expected to equal the mutation rate, as population size is cancelled out (Kimura 1968). Since a migration event effectively mirrors a mutation event, as both events introduce new alleles to the gene pool, a similar argument can be made with respect to the proportion of introgressed alleles within a population.

For any admixed population, the total proportion of introgressed alleles within genomes, *f*, is ultimately determined by *m*, the cumulative proportion of immigrant breeders. Owing to recombination, introgressed chromosomes will gradually break up into c-loci. For each c-locus, the expected number of introgressed alleles in diploid populations immediately after a pulse introgression event equals *2·Ne·m*. For each of these introgressed alleles, the probability of eventually reaching fixation is 1/(2*N_e_*). This implies that *f* is expected to simply equal *m*:

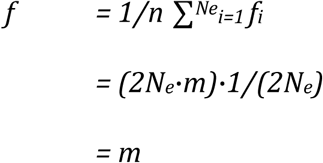

This law of conservation, *f = m,* is irrespective of the time since the gene flow event. Thus, assuming no further gene flow events, the mean proportion of introgressed alleles in genomes of an admixed population remains constant through time. What changes, instead, is the variation in the proportion of introgressed loci across individuals. Directly after the introgression event, a few hybrid individuals will carry all introgressed loci, while all other individuals are unaffected. Through generations, recombination will distribute the introgressed loci evenly throughout the population, such that eventually all individuals contain equal proportions (McFarlane and Pemberton 2019; Martin and Amos2021).

This law of conservation implies that the admixture proportion, *m*, can be derived by estimating *f*, the proportion of introgressed alleles within an admixed population. In the case of an ancient admixture event, if suffices to estimate this proportion from a single genome of that population. However, estimating *f* from a distance matrix is challenging.

Assuming that ingroup population B received gene flow from outgroup population C, which nucleotide diversity *π_c_*, the proportion of introgressed alleles is initially given by:

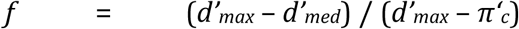

In here, *d’_max_*, *d’_med_* and *π‘_c_* refer to distances directly after the admixture event. The numerator quantifies the difference in genetic distance of the two ingroup populations *A* and *B* relative to the outgroup population *C*, a difference which is caused by the admixture event. The denominator quantifies the potential upper limit of this value, which is reached when all individuals of population *B* have been replaced with individuals of population *C* (i.e., *f* = 1). In this extreme scenario, the mean genetic distance between *B* and *C* decreases to *π’_c_*, the nucleotide diversity of donor population *C* at the time of the admixture event.

Unfortunately, *d’_max_*, *d’_med_* and *π‘_c_* cannot be simply replaced with values *d_max_*, *d_med_* and *π_c_*, observed at a certain time, *T’*, after the admixture event. This is because *π_c_* will not increase over time proportional with *d_max_* and *d_med_*, which causes the ratio between the numerator and denominator to vary.

A rare opportunity for estimating *f*, and thereby *m*, arises in the case that the outgroup population *C* happened to split into two sister populations shortly after the admixture event. Given this demographic scenario, *π‘_c_* can be replaced by the *D_xy_*-estimate for these two sister populations (*d_out_*):

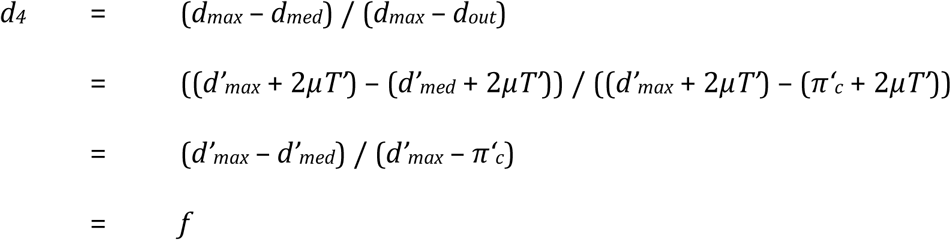

Considering that population size fluctuations can cause a rapid decrease but not increase of nucleotide diversity, it is likely that *π_c_* was at a higher level during the admixture event than at the time of the subsequent population split. Hence, the *d_4_*-score may be considered a conservative estimate of the true proportion of introgressed alleles.

### BOX 1. Numerical example of calculating *E(p)* for a pair of diploid individuals

Consider two diploid individuals, *x* and *y*, with the following haplotypes for a 5-bp locus:

| *ind* | *allele* | *haplotype* |
| --- | --- | --- |
| $x$ | 1 | ACGCA |
|  | 2 | TGGCA |
| $y$ | 1 | TGGTA |
|  | 2 | AGGTC |

From this phased dataset, we can infer the expected sequence dissimilarity, *E(p),* between individuals *x* and *y*, as the mean of the four possible haplotype comparisons:

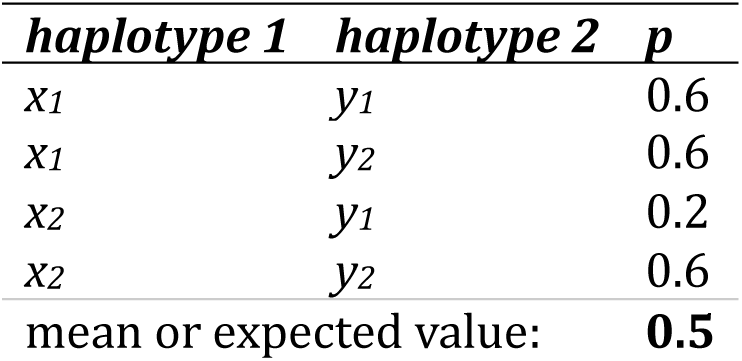

| *haplotype 1* | *haplotype 2* | *p* |
| --- | --- | --- |
| $x_1$ | $y_1$ | 0.6 |
| $x_1$ | $y_2$ | 0.6 |
| $x_2$ | $y_1$ | 0.2 |
| $x_2$ | $y_2$ | 0.6 |
| mean or expected value: |  | **0.5** |

In case the data is unphased, we cannot apply the above approach. Instead, we calculate the mean distance from single base pair distances. Per base pair, we assume the distance is 0 if individuals are homozygous for the same allele (e.g., AA and AA), 1 if individuals have no allele in common (e.g., AA and TT, or AT and CG), and 0.5 if one or both individuals are heterozygous and have at least one allele in common (e.g. AT and TT, or AT and AT):

| *site* | *alleles* |  | *genotypes* |  | *p* | *ASD* | *Hamming* |
| --- | --- | --- | --- | --- | --- | --- | --- |
|  | *x* | *y* | *x* | *y* |  |  |  |
| 1 | AT | TA | 1 | 1 | 0.5 | 1 | 0 |
| 2 | CG | GG | 1 | 0 | 0.5 | 1 | 1 |
| 3 | GG | GG | 0 | 0 | 0 | 0 | 0 |
| 4 | CC | TT | 0 | 2 | 1 | 2 | 1 |
| 5 | AA | AC | 0 | 1 | 0.5 | 1 | 1 |
| mean: |  |  |  |  | **0.5** | 1.0 | 0.6 |

By estimating sequence dissimilarity, we arrive at the same conclusion, namely E(p)=0.5. For comparison, we included sharing distance (ASD) and Hamming’s (bitwise) distances.

### BOX 2. Numerical example of calculating *He*, *π*, *D_xy_*, *D_a_* and *F_ST_* from *E(p)*

Consider the following dataset of multi-locus *E(p)*-estimates between four diploid individuals, which belong either to population *A* (A1, A2) or population *B* (B1, B2):

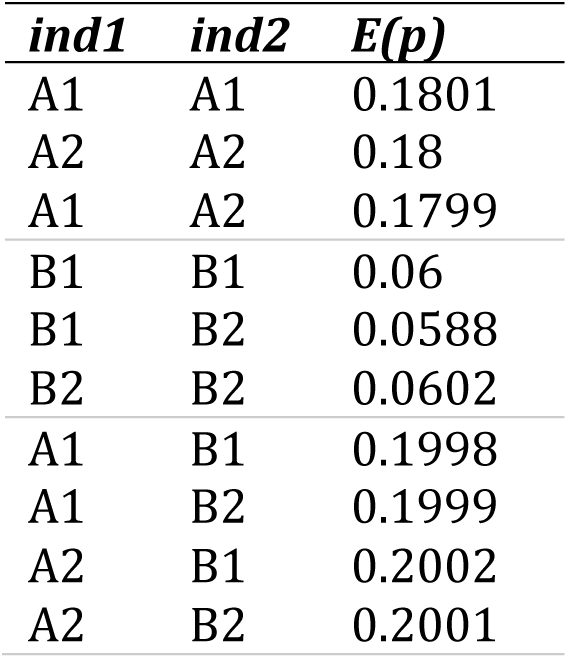

For within-individual comparisons, *E(p)*-estimates denote heterozygosity. By taking the mean of all within-population comparisons, we can infer the nucleotide diversity of the two populations, namely: *π_a_* = 0.18, and *π_b_* = 0.06. By taking the mean of between-population comparisons, we can infer the absolute genetic distance between the two populations: *D_xy_* = 0.2.

Mean nucleotide diversity of the two populations equals: *π_xy_* = (0.18 + 0.06)/2 = 0.12. With this *π_xy_*-estimate we can subsequently infer net divergence and relative genetic distance using the formula’s: *D_a_* = *D_xy_* – *π_xy_* and *F_ST_* = *D_a_*/*D_xy_*. We obtain: *D_a_* = 0.2 – 0.12 = 0.08, and *F_ST_* = 0.08/0.2 = 0.4.

This *F_ST_*-value of 0.4 indicates that the average pairwise sequence dissimilarity within populations is 40% smaller than the sequence dissimilarity between populations (i.e., 10% for population *A*, and 70% for population *B*) (M. J. de Jong et al. 2024).

### Box 3. Numerical example of calculating *d3*- and *d4*-scores

Consider the following distance matrix with *D_xy_*-estimates between three populations:

|  | A | B | C |
| --- | --- | --- | --- |
| A |  | 0.14 | 0.2 |
| B |  |  | 0.136 |
| C |  |  |  |

If assuming a strict molecular clock, we can test for admixture by calculating the *d3*-score:

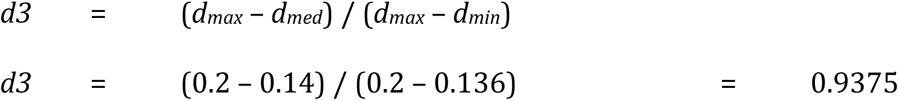

This value can range between 0 and 1, with 0 indicating no admixture and 1 indicating that the intermediate population is equally distant to the remaining two populations.

One of the many potential underlying scenarios, is that population B has a hybrid origin, and received 60% of its founder coming from population A and 40% coming from population C. This would result in the observed distance matrix if C had a nucleotide diversity of 0.04, and if the *D_AB_* was originally 0.1.

If assuming that directly after the admixture event, population C split into subpopulations C1 and C2, and that no further admixture events occurred, the admixture proportion can be determined at any moment in time using the *d4*-score. For instance, assuming novel mutations added 0.1 to all between-population distances:

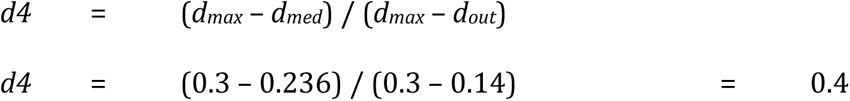

## CONCLUSIONS

We present DIST, a distance-based method for species tree inference. The key rationale underlying DIST is that the tree of individuals approximates an imaginary average gene tree. All branch lengths of this average gene tree reflect mean coalescence time, *E(t)*, which depend on population split times (*T*) and effective population sizes (*Ne*), and hence are accurately summarised by a species tree constructed from an *F_ST_*-distance matrix. DIST generates this distance matrix by calculating mean values of sequence dissimilarity within and between populations, known as *π* and *D_xy_*, and by subsequently inferring Hudson *F_ST_*-estimates. Given certain evolutionary scenarios, such as absence of novel mutations or population size constancy, branch lengths can be converted from *F_ST_*-estimates into coalescent units (*τ*), such that the species tree becomes an explanatory model. Optionally, the species tree can be transformed into a reticulated network by adding gene flow edges, in order to account for reticulate evolution. Gene flow events are detected by testing for observed deviations from equidistance of an outgroup population to two ingroup populations. DIST quantifies this deviation using the *d3*-score, and applies this method to all possible population triplets.

## Availability

A bash-script to calculate *E(p)*-estimates from an input gVCF-file can be obtained from Github: https://github.com/mennodejong1986/DIST. The resulting data frame can be used as input for distance-based phylogenetic inferences with the R package SambaR: https://github.com/mennodejong1986/SambaR.

---

**Supplementary Table S1.**
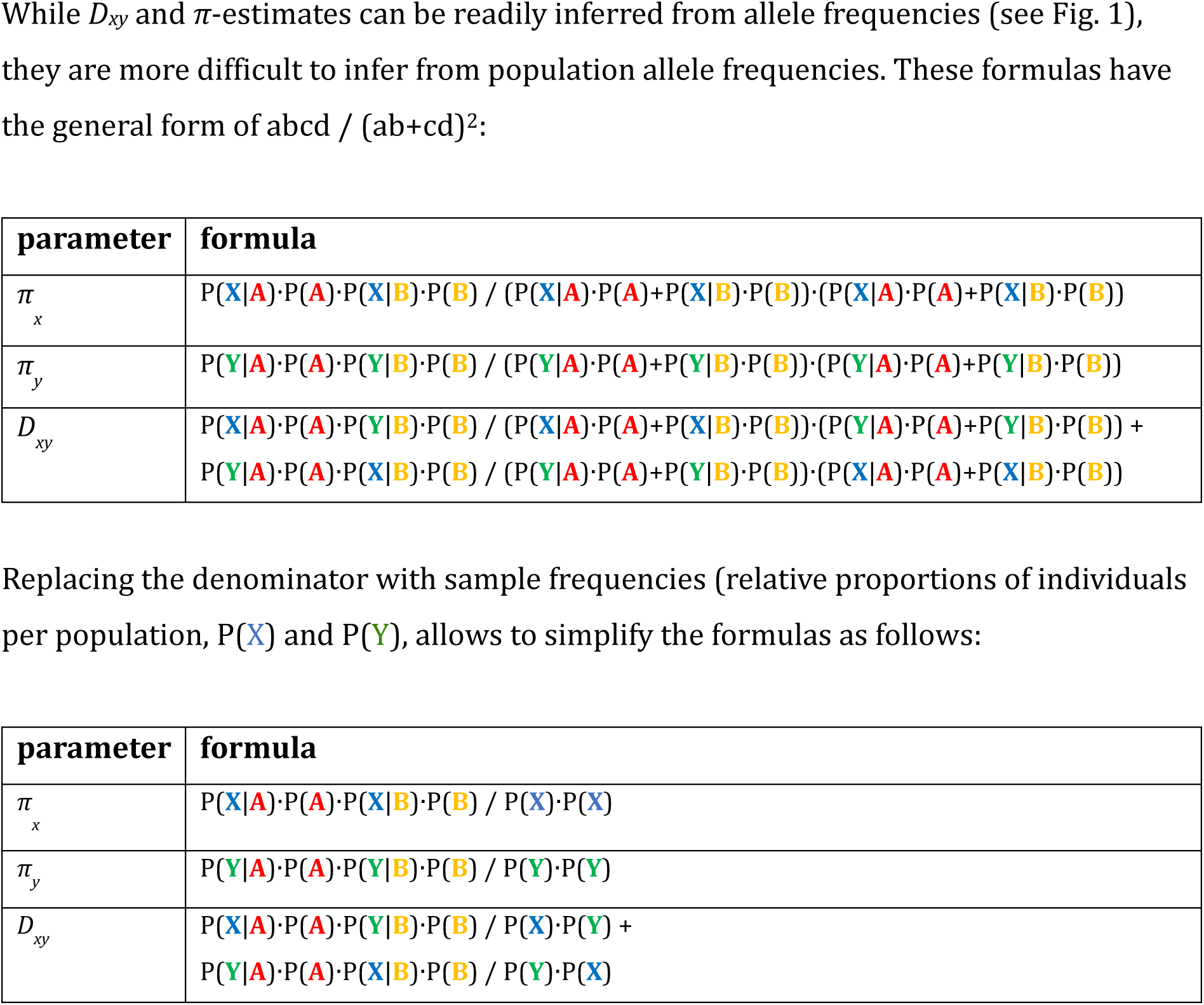

**Figure S1.**
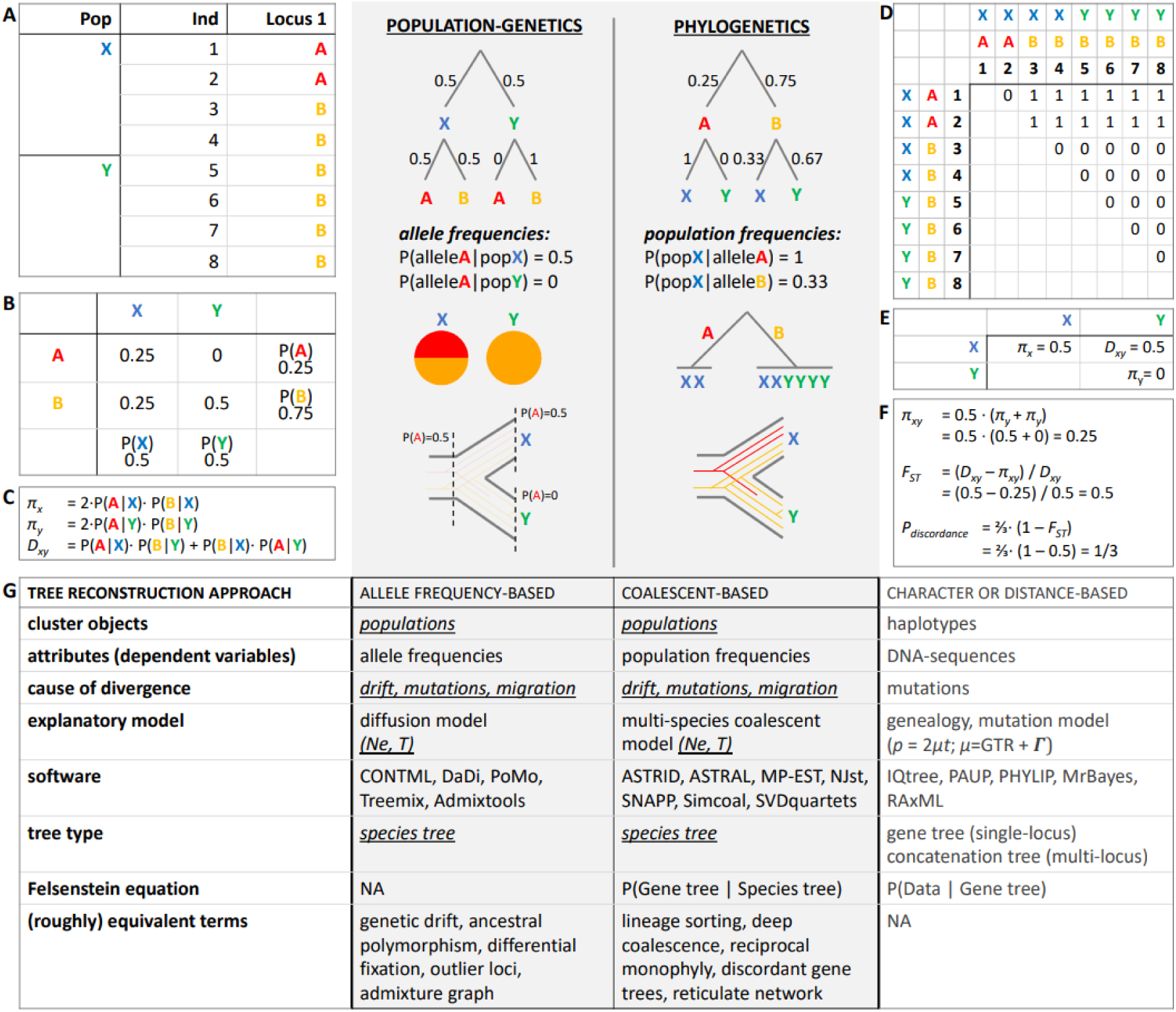
Two sides of the same coin: population-genetics versus phylogenetics. Whereas population-genetics categorise for each locus the observed genetic variation among individuals by population, phylogenetics do so by allele (A-B). This maintains two parallel research fields which target the same objective, species tree inference, using very different approaches (G). Distance-based approaches side-step this fundamental design choice by calculating distances between each pair of individuals (D). Population-level distances, *D_xy_* and *π*-estimates, which can be readily inferred from the distance matrix as well as from allele frequencies (C, E), can be used to quantify the expected degree of gene tree discordance (F), i.e., to quantify how often a haplotype from population X is more similar to a haplotype of population Y than to the other haplotype of population X.

**Figure S2.**
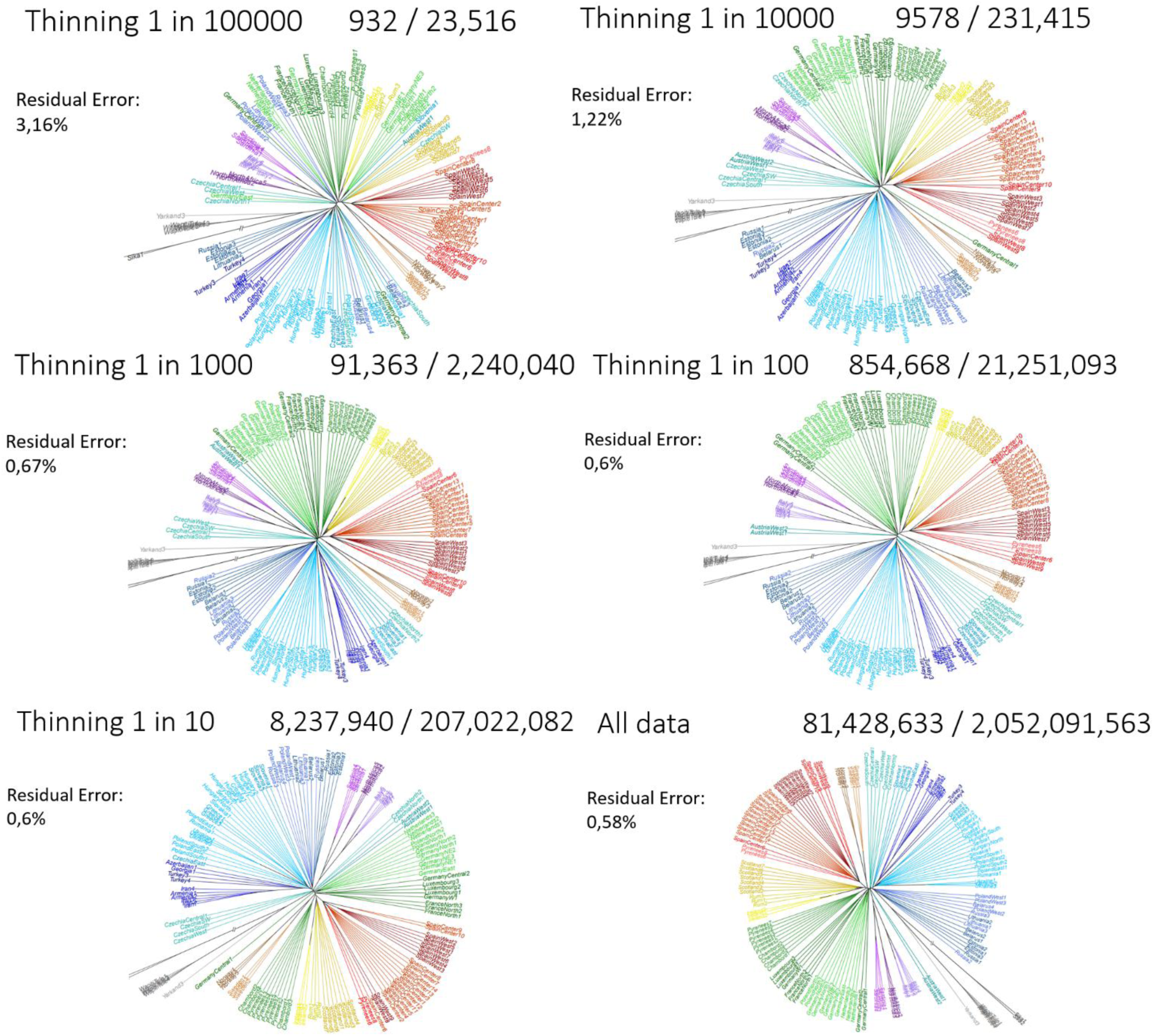
Sensitivity to thinning factor. Distance-based concatenation trees (‘trees of individuals’) for a dataset of *Cervus elaphus*, from De Jong et al. (2025). Note that increased resolution leads to roughly ultrametric trees, while the topology remains stable for thinning parameters up to 1 in 1000. The numbers before and after the forward slash indicates the number of variable sites and number of total sites, respectively. For instance, the tree of individuals for the unthinned dataset was inferred from >81M variable sites, out of >2Gb sites in total.

